# Female-Biased IL-17–STAT3 Signaling Marks Dementia-Associated Microglial Inflammation in Alzheimer’s Disease

**DOI:** 10.64898/2026.09.16.752221

**Authors:** Nourhan Abdelfattah, Reece Kang, Dante Jimenez, Sonia Villapol, Juan Toledo, Timothy Hohman, Sung Yun Jung, Kyuson Yun

**Affiliations:** Department of Neurology, Houston Methodist Research Institute, Houston, TX 77030, USA; Department of Neurosurgery, Houston Methodist Research Institute, Houston, TX 77030, USA; Vanderbilt Memory and Alzheimer’s Center, Vanderbilt University Medical Center, Nashville, TN 37203, USA; Department of Biochemistry, Baylor College of Medicine, Houston TX; Department of Neurology, Weill Cornell Medical College, New York, NY 10065, USA

**Keywords:** Alzheimer’s disease, sex difference, microglia, neuroinflammation, STAT3, dementia, signaling pathway, single cell analysis, IL17

## Abstract

**INTRODUCTION:** Alzheimer’s disease (AD) disproportionately affects women, but the underlying mechanisms are not fully understood. We compare sex-specific molecular changes across AD with and without dementia at single-cell resolution.

**METHODS:** We performed sex-stratified single-nucleus transcriptomic analysis of human AD to dissect sex-specific differences in dementia progression, using the SEA-AD cohort for discovery and the ROSMAP and NBB datasets, together with 5xFAD mouse models, for validation. Donors were classified as no-dementia controls, no-dementia AD, or dementia AD, followed by donor-level pathway analysis, de novo microglial clustering, communication modeling, and cytokine-response enrichment

**RESULTS:** Females with AD dementia showed enhanced inflammatory pathway activity across glial, neuronal and vascular-associated cell types, with the strongest enrichment in microglia. This female-biased program centered on IL6-JAK-STAT3 signaling and increased STAT3 expression in microglia and astrocytes. These results were validated in the ROSMAP and NBB datasets and conserved in the 5xFAD mouse model. Microglial clustering resolved 18 states and revealed sex-divergent disease remodeling, including female-specific expansion of redox-reactive and CD163+ macrophage-primed states. Cell:cell communication analysis identified female-biased TGFβ/GRN signaling, male-biased CD99 signaling, and dementia-associated SPP1-integrin communication with the vasculature. Microglial cytokine-response analysis identified IL-2 response as a dementia-associated signature in both sexes, alongside a female-biased IL-17/interferon response and a male-biased IL-7 response.

**DISCUSSION:** These findings identify a disease-emergent, female-biased STAT3-centered neuroinflammatory program in human and mouse AD, linking sex-specific microglial remodeling to cytokine responsiveness and glial-vascular communication. The results suggest that sex-specific changes in neuroinflammation pathways in microglia and other cell types are associated with progression to dementia.

## 1. BACKGROUND

Alzheimer’s disease (AD) is the predominant cause of age-related dementia worldwide and is anticipated to affect 50 million people by 2030, with women accounting for approximately two-thirds of all AD cases globally^1^. This sex-biased incidence was historically ascribed to female longevity, but accumulating evidence from age-stratified analyses refutes this simple explanation; women face elevated age-specific incidence rates of AD relative to men, particularly at older ages, and experience faster cognitive decline, greater neuropathological burden at autopsy, and accelerated brain atrophy following diagnosis^2,3^. Female APOE4 carriers face compounded risk, exhibiting higher cerebrospinal fluid tau levels, worse memory performance, and an earlier age of clinical onset compared to male carriers^4–6^. Despite carrying greater pathological burden, women also demonstrate paradoxically higher cognitive reserve, requiring a heavier tau load before clinical manifestation of dementia, a resilience advantage that is ultimately overwhelmed by a more aggressive disease trajectory^7–9^. These observations collectively indicate that biological sex is not merely an epidemiological covariate but a fundamental biological variable that shapes the molecular pathogenesis, cellular vulnerability, and clinical progression of AD.

At the genomic level, sex-specific genetic architecture has been identified across several stages of AD pathogenesis including sex-interacting loci associated with both CSF Aβ42 and tau^10^, a stronger APOE-ε4 to CSF tau association in women independent of amyloid or tangle burden^4^, and sex-specific genetic drivers of cognitive resilience to AD neuropathology^11^. Together, these findings point to distinct, sex-dependent genetic contributions along the amyloid to tau to cognitive outcome cascade. Multiple studies have also characterized sex differences in AD using bulk transcriptomics approaches, implicating microglia and immune-related gene networks in sex-specific AD pathology, including a female-specific network driver (LRP10) with downstream effects on neuronal and microglial population^12,13^. Cross-species analysis using mouse models has reinforced the female bias in AD neuroinflammation. In the 5xFAD and 3xTg models, female mice accumulate higher Aß42 burdens, develop spatial memory deficits earlier, and exhibit more pronounced neuroinflammation relative to male littermates^14–16^. These efforts shift our understanding of sex-differences in AD pathology to the sexually dimorphic brain, a dynamic cellular ecosystem comprising neurons, astrocytes, microglia, oligodendrocytes, and vascular cells whose coordinated interactions increasingly appears to drive neurodegeneration^17^. Single-nucleus RNA sequencing (snRNA-seq) has been transformative in this regard, enabling transcriptome-wide characterization of cell-type-specific disease states at population scale from archival frozen tissues^18^. Large-scale snRNA-seq profiling of postmortem human brain tissue has revealed that microglia and astrocytes undergo profound transcriptional reprogramming in AD, transitioning from homeostatic to pathologically reactive states that actively modulate amyloid clearance, tau propagation, synaptic integrity, and intercellular signaling networks^19–22^. Additionally, transcriptome-wide association analysis leveraging the ROSMAP snRNA-seq dataset identified 2,660 sex-specific gene-endophenotype associations, with female-protective associations enriched predominantly in neurons and female-risk associations concentrated in glial populations^22^. The vascular compartment is also linked to AD pathology: endothelial cells carry sex-biased transcriptomes reflecting differential hormonal exposures across the lifespan, and vascular contributions to AD pathology manifest in distinct ways between men and women^22–24^.

A critical gap remains in understanding how sex-specific cellular changes map onto the sex-specific clinical expression of AD. Here, we leverage human snRNA-seq datasets and mouse spatial transcriptomic data to identify a disease-emergent, sex-biased modulatory programs across glial and vascular-associated compartments, with a particular focus on the transition from AD neuropathology without dementia to AD-associated dementia. By comparing transcriptional landscapes across no-dementia controls, donors with AD neuropathology without dementia, and donors with AD-associated dementia in a sex-stratified framework, we identify coordinated inflammatory programs in female microglia, astrocytes, and vascular-associated cells that are absent or attenuated in males at comparable disease stages.

## 2. METHODS

### 2.1 Meta-analysis of the SEA-AD, ROSMAP and NBB datasets

We leveraged the publicly available snRNA-seq dataset from 84 donors in the SEA-AD^25^ (Seattle Alzheimer’s Disease Brain Cell Atlas) (AD Knowledge Portal Accession Number: syn52146347) which consisted of 1.2 million nuclei derived from 84 donors’ postmortem dorsolateral prefrontal cortex (DLPFC) (Brodmann area 9) tissues. The pre-aggregated h5ad object was downloaded and re-analyzed using Seurat (v5.2.1; RRID:SCR_016341). We also used snRNA-seq data from DLPFC brain specimens derived from 465 ROSMAP^26^ participants (AD Knowledge Portal Accession Number: syn31512863). The ROSMAP datasets contained 424 unique participants with a 1.64 million nuclei in total. The original object containing preprocessed microglia cells was downloaded and used as a validation dataset. Finally, we used the NBB (Netherlands Brain Bank) (GSE260461) ^27^ as a second independent validation cohort.

### 2.2 Cohort definitions and clinicopathological classification

To enable direct comparison of sex-specific effects across independent datasets, donors in each cohort were stratified using cognitive status at last assessment and AD neuropathologic change. Cognitive status was classified as no dementia or dementia, and AD neuropathologic change was collapsed into no/low versus intermediate/high pathology based on established neuropathological staging (**Figure 1A**). Crossing these axes defined four groups: NoDementia.Control, NoDementia.AD, Dementia.AD, and dementia without AD pathology **(Supplementary Figure 1A**). The dementia without AD pathology group was excluded from downstream analyses so that Dementia.AD refers to clinically and pathologically confirmed AD-associated dementia rather than dementia of any etiology.

**Figure 1:**
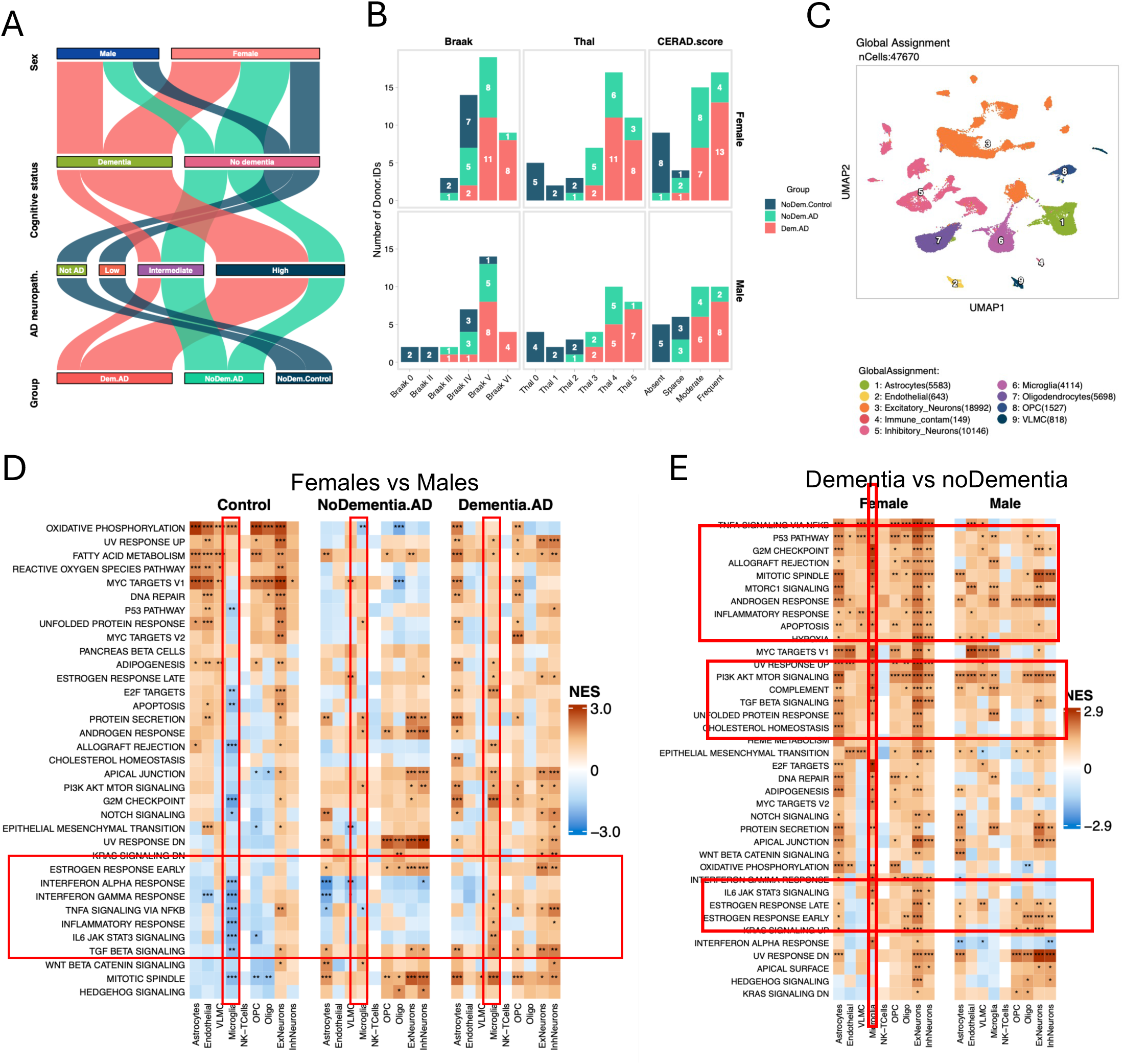
Disease stage and sex specific molecular changes in human AD. **A.** Parallel sets (alluvial) diagram showing sex, cognitive status, and neuropathology criteria considered for assigning each donor to a comparison group. Stream width is proportional to donor count. Streams are colored by Group level. Middle axes: Overall.AD.neuropathological.Change, Cognitive.Status. **B.** Stacked bar chart summarizing the distribution of categorical metadata variables deduplicated to one row per Donor.ID. Each column panel corresponds to a separate metadata variable (Braak, APOE.Genotype, CERAD.score). Bars are stacked and colored by Dementia.AD, with each segment labeled with its count. Rows are split by Sex.**C.** Single-nucleus landscape of the SEA-AD cohort. UMAP of 1,598,085 nuclei from 76 donors, integrated with Harmony across donor identity,library prep method; Louvain clustering (resolution 0.5) resolved 32 clusters annotated to 9 global cell types. Also see Supplementary Figure 1. **D.** Heatmap of pseudobulk (Σ raw counts per donor × cell type → DESeq2) GSEA Normalized Enrichment Scores (Hallmark) for differentially enriched in female vs male samples, analyzed separately by disease stage. in Control, NoDementia.AD or Dementia.AD across cell types. Rows = union of top 20 gene sets by max |NES| across cell types. **E.** Heatmap of pseudobulk GSEA Normalized Enrichment Scores (Hallmark) for differentially enriched in Dementia vs NoDementia samples, analyzed separately by sex across cell types. Rows = union of top 30 gene sets by max |NES| across cell types. Method: donor-level pseudobulk, edgeR TMM normalization, limma-voom moderated t-statistic pre-ranking, fgsea. NES > 0 = enriched in the first term of the contrast. Overlaid significance stars: * BH < 0.05, ** < 0.01, *** < 0.001.

### 2.3 SEA-AD cohort

Donor-level cognitive status and neuropathologic change were obtained from the SEA-AD^25^cohort metadata. ADNC was assigned using the NIA-AA “ABC” consensus criteria^28^, which combine Thal amyloid phase (A), Braak neurofibrillary tangle stage (B), and CERAD neuritic plaque score (C) into a four-level classification (Not AD, Low, Intermediate, High). Donors with Intermediate or High ADNC were classified as AD-pathology-positive; those with Not AD or Low ADNC were classified as pathology-negative. Crossing this with clinical cognitive status (Cognitive Status: No dementia vs. Dementia) yielded the four groups described above. Three donors without cognitive or neuropathological assessment (young reference brains) were excluded prior to classification, and four donors with dementia but without AD pathology (non-AD dementia) were also excluded from the analysis cohort. The final retained cohort comprised 76 donors: NoDementia.Control (n=17; 9 female, 8 male), NoDementia.AD (n=24; 15 female, 9 male), and Dementia.AD (n=35; 21 female, 14 male).

### 2.4 ROSMAP cohort

Donor clinical and neuropathological data were obtained from the Religious Orders Study and Memory and Aging Project (ROSMAP). Cognitive status was derived from the last-visit clinical dementia diagnosis (dcfdx_lv), established through the ROSMAP protocol’s three-stage process (computerized scoring of a 19-test cognitive battery, review by a neuropsychologist, and final diagnostic classification by a clinician) per NINCDS/ADRDA criteria. Per the ROSMAP clinical codebook, donors with no cognitive impairment or with mild cognitive impairment (with or without a contributing comorbid condition) were classified as No dementia; donors with a clinical diagnosis of Alzheimer’s dementia (with or without a contributing comorbid condition) or of another primary dementia were classified as Dementia. ADNC was approximated from Braak neurofibrillary tangle stage and CERAD neuritic plaque score, following the same A×B×C framework as above; Thal amyloid phase was not available for the majority of ROSMAP donors in this data release and was therefore not incorporated as an input to ADNC in this cohort, a methodological difference from the SEA-AD classification that should be considered when interpreting cross-cohort comparisons. Donors with a clinical diagnosis of dementia attributable to a cause other than Alzheimer’s disease were classified as non-AD dementia regardless of incidental AD pathology and excluded from the analysis cohort, consistent with the SEA-AD design. The final retained cohort comprised 403 donors: NoDementia.Control (n=140; 88 female, 52 male), NoDementia.AD (n=127; 87 female, 40 male), and Dementia.AD (n=136; 104 female, 32 male).

### 2.5 NBB cohort

Donor-level clinical and neuropathological data were obtained from the donor set associated with Rosenzweig *et al*^27^. Clinical diagnosis (Non-demented control vs. Alzheimer’s disease) was established by the NBB through antemortem medical record review, and neuropathological status was assessed using Braak neurofibrillary tangle stage and CERAD neuritic plaque score. Since clinical annotation of dementia was not available for this dataset^27^, donors were stratified into Control and AD groups only, without further subdivision by cognitive-pathological concordance, consistent with the original study design. The final cohort comprised 36 donors stratified by sex: Control (n=18; 10 female, 8 male) and AD (n=18; 8 female, 10 male).

### 2.6 Donor Levels Gene Set Enrichment Analyses (GSEA)

Donor-level gene set enrichment analysis was performed using a pseudobulk framework. Within each major cell type, raw RNA counts were summed for each donor to generate one donor-level expression profile per cell type. Donor-by-cell-type combinations with fewer than 10 nuclei were excluded, and contrasts were tested only for groups with at least three donor samples. Pseudobulk libraries were normalized using trimmed mean of M-values normalization in edgeR, and mean-variance relationships were modeled using limma-voom. Linear models were fit using a group-means design, with APOE genotype and age at death included as covariates. Genes were ranked by moderated t-statistics from limma, and pathway enrichment was assessed using fgsea against the MSigDB Hallmark collection. A positive normalized enrichment score indicates enrichment in the first group named in each contrast. Pathways with a maximum absolute normalized enrichment score below 1 across all contrasts were omitted from summary heatmaps.

### 2.7 Cellular composition analysis

Per-donor fractions of each cell type (of total nuclei) were computed from cluster assignments. Compositional differences were modeled with a quasibinomial generalized linear model (fraction ∼ Disease stage × Sex + APOE ε4 + Age + PMI); disease and sex-by-disease interaction effects were tested by nested-model likelihood-ratio (F) tests, with BH correction applied across cell types. Pairwise group comparisons (t-test, BH-adjusted) are shown for visualization alongside the omnibus model.

#### Cell-cell communication analysis

Intercellular communication was inferred with CellChat^29^(v2.2.0, RRID: SCR_021946), run independently for each sex-by-disease-stage groups, with nuclei downsampled to 2,000 per global cell type within each group (total 82,433 cells across 6 conditions (Female.Dementia.AD; Female.NoDementia.AD; Female.NoDementia.Control; Male.Dementia.AD; Male.NoDementia.AD; Male.NoDementia.Control) to control for cell-number-dependent bias in communication-probability estimation. Signaling pathways were ranked by the coefficient of variation and range of total information flow (summed communication probability across all sender-receiver pairs) across the six groups. For donor-level analyses, ligand-receptor communication probabilities were computed per donor, zero-filled for donor-link combinations in which both sender and receiver cell types were present at sufficient abundance but no communication was detected and compared between sexes within each disease stage (Wilcoxon rank-sum test, BH-corrected within stage) and across the disease trajectory within each sex. Communication patterns were identified for k = 3 to 10 latent patterns using non-negative matrix factorization. All analyses were conducted in R using the scSidekick^30^ package wrapper (v1.1)

### 2.8 De novo microglial clustering and consensus gene programs

Microglial nuclei were re-clustered independently of the global cell-type assignment, yielding 18 transcriptionally distinct states, annotated using each cluster’s top differentially expressed marker genes. Because baseline microglial composition differed by sex, disease-associated changes in state proportion were tested within each sex separately across the disease trajectory (pairwise Wilcoxon rank-sum tests on per-donor state fractions: Control vs. NoDementia.AD, NoDementia.AD vs. Dementia.AD, Control vs. Dementia.AD), rather than by comparing sexes at a fixed disease stage. Consensus non-negative matrix factorization (cNMF) was applied to the microglial expression matrix to recover shared transcriptional programs independent of discrete cluster boundaries (14 programs). cNMF detection and integration was done in R using the scSidekick^30^ package wrapper (v1.1, RRID:SCR_028835).

### 2.9 Upstream cytokine-response enrichment analysis (Parse Biosciences cytokine atlas)

To infer upstream cytokines associated with sex-biased microglial programs, we used the publicly available human PBMC cytokine atlas^31^, in which peripheral blood mononuclear cells from 12 donors were stimulated with 90 recombinant cytokines and profiled by single-cell RNA sequencing to catalogue cytokine induced gene expression changes in PBMCs. We downloaded the precomputed differential expression tables released with the dataset. For each cytokine, we restricted analysis to the CD14+ monocyte population, the myeloid cell type most transcriptionally comparable to microglia in the atlas, and defined a cytokine-response signature from the top up-regulated genes relative to unstimulated controls. This produced a custom collection of 87 cytokine-response gene sets. Enrichment of each cytokine-response signature in the discovery cohort was quantified with the scSidekick^30^ GSEA tools (v1.1, RRID:SCR_028835). Counts were aggregated to donor-level pseudobulk (summed raw counts per donor, or per donor and cell type) and log-CPM normalized. Single-sample GSEA (ssGSEA) scores were computed for every cytokine signature in each donor using GSVA::gsva() (method = “ssgsea”). Donor-level ssGSEA scores were then compared across the six sex-by-stage groups (female and male; NoDementia.Control, NoDementia.AD, Dementia.AD). To test which cytokine signatures are engaged during the disease transition, we performed donor-level pseudobulk GSEA against the same cytokine gene sets. Genes were normalized with edgeR TMM, modeled with limma-voom, and pre-ranked by the moderated t-statistic; enrichment was computed with fgsea. We evaluated the contrasts Dementia.AD versus NoDementia.AD and Dementia.AD versus NoDementia.Control, separately within female and male donors. Normalized Enrichment Scores (NES) are reported with Benjamini-Hochberg-adjusted significance (* BH < 0.05, ** < 0.01, *** < 0.001). For the cell-resolved analyses (butterfly plot), each microglial cell (n = 38,016) was scored against four signatures (IL-7, 50 genes; IL-17D, 50 genes; TWEAK, 20 genes; IL-17F, 2 genes) as the mean expression of the signature genes, column-centered across cells. A two-axis hierarchy was constructed in which the y-axis separates the IL-7/IL-17D programs (y > 0) from the TWEAK/IL-17F programs (y < 0) and the x-axis encodes the log2-scaled left-right balance within the dominant axis. Cells were visualized in aggregate and split by group using the scSidekick^30^ package wrapper (v1.1, RRID:SCR_028835).

### 2.10 Animal studies

Male and female 5xFAD hemizygous mice (C57BL/6J; Jackson Laboratory, 034848) and wild-type littermates (Jackson Laboratory, 000664) were age-and sex-matched at the time of all experiments. Mice were randomly assigned to experimental groups. No mice were excluded from the survival study. All procedures were approved by the Houston Methodist Research Institute Institutional Animal Care and Use Committee

### 2.11 Immunofluorescence validation

Amyloid-β clone 6E10 (BioLegend Cat# 803004, RRID:AB_2715854), IBA1 (Thermo Fisher Scientific Cat# PA5-27436, RRID:AB_2544912), pSTAT3 (Cell Signaling Technology Cat# 9145, RRID:AB_2491009),STAT3 (Cell Signaling Technology Cat# 9139, RRID:AB_331757) and SPP1 (R&D Systems Cat# AF808, RRID:AB_2194992) protein expression were quantified by immunofluorescence in 5xFAD mice, comparing female and male animals.

### 2.12 5XFAD mouse Visium dataset

Spatial transcriptomic data were obtained from the published 10x Genomics Visium dataset GSE233208 (5xFAD mouse model-Miyoshi *et al.*), supplied as a processed and annotated Seurat object containing 212,249 spots across 80 capture areas, with per-spot region annotations already assigned by the original authors^32^. This dataset profiled 5xFAD and wild-type mouse brains across 4-12 months of age. Each capture area corresponds to one mouse/one tissue section on one slide. Spot-level scores were averaged within section/mouse before any test, so each section contributes exactly one observation to the whole-brain contrast and the number of spots does not inflate significance. Stratified analyses average within section and group. Canonical STAT3 targets and regulators (30 genes, *A2m, C3, Ccnd1, Cd14, Cebpb, Cebpd, Cp, Fos, Gfap, Hspb1, Il10ra, Il11ra1, Il6ra, Il6st, Irf1, Jak1, Jak2, Junb, Lifr, Mcl1, Myc, Nfkbia, Osmr, Ptpn11, Serpina3n, Socs3, Stat1, Stat3, Timp1, Vim*) Keren-Shaul disease-associated microglia ^33^ (DAM: *Aif1, Apoe, Axl, B2m, Ccl6, Cd63, Cd68, Cd9, Csf1, Cst7, Ctsb, Ctsd, Ctss, Ctsz, Cx3cr1, Fth1, Gpnmb, Hexb, Igf1, Itgax, Lgals3, Lgals3bp, Lpl, P2ry12, Spp1, Timp2, Tmem119, Trem2, Tyrobp*) signature modules were scored per spot with UCell (v2.14.0) (‘ScoreSignatures_UCell’) on raw counts with ‘maxRank = 1500’. UCell is rank-based within each spot, which makes scores robust to differences in total counts between spots but means scores are comparable *between groups for the same module*, not between different modules in absolute magnitude.

### 2.13 Plaque-proximity proxy

Fourteen plaque-associated genes were selected for a 5xFAD/WT expression ratio > 2 and detection in at least 2% of spots: *Cst7, Itgax, Ccl6, Gpnmb, Lyz2, Trem2, Cd68, Mpeg1, Tyrobp, Ctsz, Ctss, Lpl, Cd9,* and *Gusb*. We also calculated PIG_plaque scores^34^ using *Cst7, Ctss, Ctsb, Ctsd, Ctsz, Cd68, Cd9, Itgax, Tyrobp, Trem2, Apoe, Clu, C1qa, C1qb, C1qc, C4b, Gfap, Vim, Serpina3n, Hexb, Laptm5, Fth1, Ftl1, Lyz2, B2m, Cd52, Npc2, Grn, Lgals3bp,* and *Mpeg1*. The list was scored per spot with UCell under the same settings, then smoothed over the Visium hexagonal lattice by averaging each spot with itself and its six immediate neighbors, identified within each section image using array-coordinate offsets. Section-level burden was defined as the mean smoothed score across all spots in that section.

### 2.14 Image-based amyloid plaque quantification

The transcriptional proxy above establishes association but not amyloid burden itself. To measure plaque load directly, the H&E/immunofluorescence section images stored in the Seurat object were analyzed independently of the transcriptome. Only 64/80 sections contained amylo glo stained plaques (same as original manusctipt^32^). Plaque-like puncta were detected on the magenta channel, defined per pixel as ‘min(R, B)’, which is high where the stain is present and low for achromatic debris. All computation was confined to a tissue mask constructed from the Visium spot coordinates (a ±6 px box around each spot center), so that off-tissue background could not contribute. Plaque-like puncta were then visualized and confirmed per section.

### 2.15 Partial correlation analysis

To isolate pathology-specific associations between microglial state composition and neuropathologic burden, we computed partial Spearman correlations between donor-level state composition (percentage of total microglia) and each pathology measure (Braak tau stage; CERAD amyloid score), controlling for the other pathology measure and APOE4 carrier status. For each state, sex, and pathology axis, we first residualized both the composition variable and the pathology measure of interest against the remaining covariates using linear regression, then computed the Spearman correlation between the two residual sets. This approach yields a standardized, bounded effect size (analogous to a simple correlation) that reflects the variance uniquely attributable to the pathology axis of interest, rather than variance shared with the collinear covariate, which is necessary in this cohort given the substantial correlation between amyloid and tau burden. All analyses were performed separately by sex. Formal significance for each state-sex-pathology combination was assessed independently using a quasibinomial generalized linear model (state fraction ∼ Braak + CERAD+ APOE4), with the partial correlation serving as the corresponding effect-size measure for visualization.

## 3. RESULTS

### 3.1 SEA-AD Cohort characteristics

The goal of this study is to identify sex-specific molecular changes in various cell types during AD progression in the human brain. For the discovery phase, we analyzed the publicly available SEA-AD cohort single nuclei RNA-sequencing (snRNA-seq) dataset, selected for its harmonized clinical and neuropathological metadata and its sampling across the clinical-pathological spectrum of AD. To distinguish AD neuropathology from its clinical expression as dementia, donors were classified using two independent criteria: cognitive status, defined as no dementia versus dementia, and overall AD neuropathologic changes, derived from Thal amyloid phase, Braak neurofibrillary tangle stage, and CERAD neuritic plaque score using the NIA-AA ABC criteria and collapsed into no/low versus intermediate/high AD pathology (**Figure 1A-B, Supplementary Figure 1A**). Crossing these axes yielded four possible groups. The three retained groups were NoDementia.Control, donors without dementia and with no or low AD neuropathology (n = 17; 9 female, 8 male); NoDementia.AD, donors without dementia despite intermediate or high AD neuropathology, representing a pathology-positive dementia-free state at last assessment (n = 24; 15 female, 9 male); and Dementia.AD, donors with both dementia and intermediate or high AD neuropathology, representing clinically and pathologically confirmed AD dementia (n = 35; 21 female, 14 male). The fourth group, donors with dementia but no or low AD pathology, was excluded to ensure that the Dementia.AD group included only AD-associated dementia and not non-AD dementia. The final analysis cohort therefore included 76 donors, comprising 45 females and 31 males (**Figure 1A-B, Supplementary Figure 1A**).

The three comparison groups were well matched on potential confounding variables. Donors were comparable in age at death (overall 89 ± 8 years; group means 88.2 ± 9.5, 91.1 ± 5.8 and 88.0 ± 8.1 years; Kruskal-Wallis p = 0.46) and in years of education (overall 16 ± 3 years; p = 0.45), and the sex distribution (Fisher exact p = 0.90), ensuring that within-group sex comparisons are not confounded by group composition. As expected from the group definitions, Braak stage and CERAD score increased markedly along the disease trajectory (both p < 0.001), and cognitive performance declined specifically in the Dementia.AD group (last CASI 80.2 ± 8.7, MMSE 22.4 ± 4.7, MoCA 14.1 ± 6.4; all p ≤ 0.01), confirming that the groups capture distinct clinical-pathological stages (**Supplementary Table 1, Supplementary Figure 1B**). APOE genotype distribution also differed across groups (Fisher p = 0.003): the proportion of APOE ε4 carriers rose along the trajectory, from 0% of Control donors to 33% of NoDementia.AD and 43% of Dementia.AD donors (**Supplementary Table 1, Supplementary Figure C**), consistent with the established role of APOE ε4 in promoting both AD neuropathology and progression to dementia.

Integration of the retained snRNA-seq data resolved into 32 clusters that were annotated to 9 global cell types and 24 subtypes (**Figure 1C**, **1D, Supplementary Figure 1D-E**). To ask whether cellular composition itself differed across the disease states or between sexes, we modelled per-donor non-neural cell types using a quasibinomial GLM adjusted for APOE genotype and including a sex-by-disease interaction and confirmed the results with Dirichlet regression. Microglial and astrocytic fractions trended to be higher in female Dementia.AD, but no cell-type fraction differed significantly by sex, disease or APOE after Benjamini-Hochberg correction (**Supplementary Figure 1F**). Therefore, cellular compositional shifts alone do not explain the sex-and disease-specific differences described below.

### 3.2 Female-biased neuroinflammatory programs in glia and vasculature

To define sex-biased transcriptional changes in various cell types across AD progression, we compared female and male donors within each clinical-pathological stage: Control, NoDementia.AD, and Dementia.AD. To avoid potential biases from unequal nuclei counts in different samples, we performed donor-level pseudobulk GSEA within each major cell type to identify pathway-level sex differences. For each donor, raw counts from nuclei belonging to the same cell type were summed to generate one bulk-like expression profile, making the donor rather than the individual nucleus the unit of analysis. Differential pathway enrichment was then tested across sex and disease-stage contrasts using limma-voom and fgsea, with models adjusted for APOE genotype and age at death (**Figure 1D, Supplementary Figure 1G**).

In controls, sex differences were dominated by metabolic programs, including oxidative phosphorylation, fatty acid metabolism, reactive oxygen species pathways, and MYC targets, consistent with baseline sex dimorphism in cellular metabolism (**Figure 1D, Supplementary Figure 1G**). Female control microglia showed lower enrichment of immune-response pathways than male control microglia, whereas female excitatory neurons showed higher TNFα and TGFβ pathway signatures. With AD pathology, these baseline differences shift; In NoDementia.AD, female astrocytes showed increased TGFβ and reduced interferon-response signature enrichments, while the male-enriched inflammatory pathways in microglia in control samples disappeared. In the Dementia.AD group, female donors showed significant enrichment of neuroinflammatory signatures in microglia compared with male donors, including TNFα/NF-κB signaling, interferon-α and interferon-γ responses, IL6-JAK-STAT3 signaling, and complement activation (**Figure 1D; Supplementary Figure 1G**). TGFβ signaling was also enriched in microglia, astrocytes, OPCs, and neurons. In addition, DNA damage response pathways, including UV response, p53 signaling, G2/M checkpoint, and mitotic spindle, were enriched in female neurons compared with male neurons.

To ask whether increased inflammation is specifically associated with dementia rather than AD pathology alone, we compared Dementia.AD with NoDementia.AD donors within each sex. Because both groups have intermediate/high AD neuropathology, this contrast enriches for transcriptional changes associated with dementia. In females, dementia was associated with broad up-regulation of inflammatory pathways across most cell types, whereas males showed weaker and more limited changes (**Figure 1E**). Microglial IL6-JAK-STAT3 signaling again illustrated this asymmetry, increasing with dementia in females but remaining largely unchanged in males (**Figure 1E**).

### 3.3 STAT3 is a convergent, female-biased hub of dementia-associated neuroinflammation

Because microglia carried the strongest sex-and dementia-associated inflammatory signal in the global cell-type analysis, we next focused on the microglial compartment. We performed donor-level pseudobulk GSEA and tested sex, disease-stage, and sex-by-disease contrasts in models adjusted for age at death and APOE genotype (**Figure 2A**). This analysis revealed a stage-dependent reversal in microglial pathway activity. Interferon-response, hypoxia, and adipogenesis pathways were relatively male-biased in control donors, attenuated in NoDementia.AD, and became strongly female-biased in Dementia.AD, consistent with a significant sex-by-disease interaction (**Figure 2A)**. The female Dementia.AD microglial program was marked by enrichment of TNFα/NF-κB, interferon, complement, and IL6-JAK-STAT3 signaling (**Figure 2A**). To corroborate these rank-based GSEA results with an orthogonal, score-based approach, we quantified donor-level microglial pathway activity using ssGSEA/GSVA. The leading differentially scored Hallmark and Biocarta pathways were dominated by neuroinflammatory programs and recapitulated the same female-biased, disease-emergent pattern observed by GSEA. These included significant enrichment of IL6-JAK-STAT3, TGFβ, complement, and allograft rejection pathways in female Dementia.AD (**Figure 2B-C, Supplementary Figure 2A, Figure 2C-D**).

**Figure 2.**
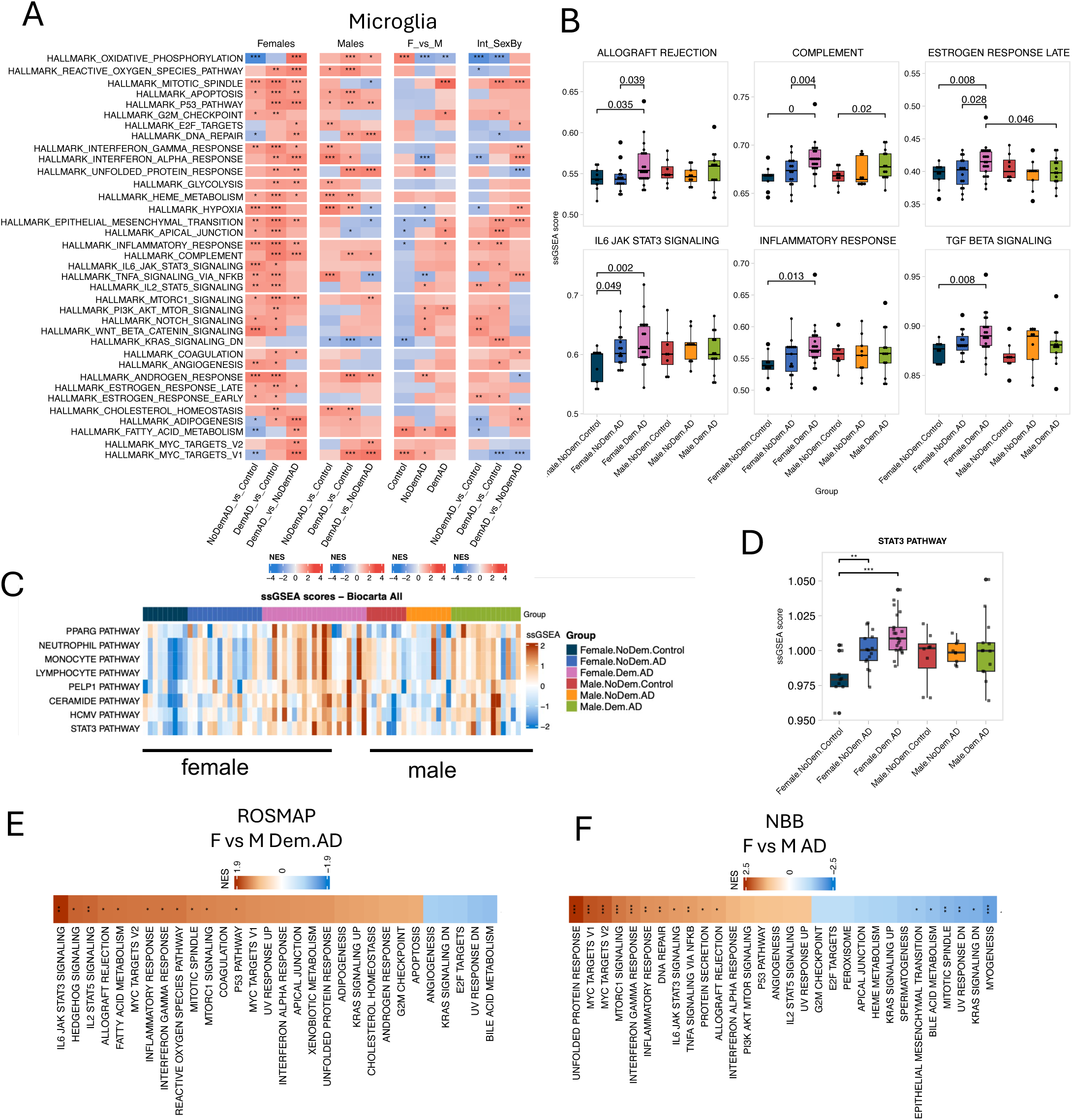
Microglia show significant female-biased neuroinflammatory programs. **A.** Summary NES heatmap for Microglia cells (Hallmark database). Rows = union of top 30 pathways per contrast, filtered to max |NES| >= 1. Columns = all contrasts tested. Stars: * pval < 0.05, ** < 0.01, *** < 0.001. **B.** Boxplots of single-sample GSEA (ssGSEA) enrichment scores for top 6 significantly differentially enriched pathways in Microglia cells (Hallmark database; BH-adjusted p < 0.05). Each panel shows score distributions per group across individual donors. **C** Heatmap and Boxplots of ssGSEA enrichment scores for top significantly differentially enriched pathways in Microglia cells (Biocarta database; BH-adjusted p < 0.05). Each panel shows score distributions per group across individual donors. **D.** Pseudobulk box plot of STAT3. Each dot represents the mean log-normalized expression for one Donor.ID (n = 76 donors), grouped by Dementia.AD, split by global cell type, rows by Sex. **E.** Pseudobulk GSEA NES heatmap (Hallmark) in ROSMAP Microglia, contrast: F vs M Dementia.AD. Stars: * BH < 0.05, ** < 0.01, *** < 0.001. **F.** Pseudobulk GSEA NES heatmap (Hallmark) in NBB Microglia, contrast: F vs M AD. Stars: * BH < 0.05, ** < 0.01, *** < 0.001.

To further test the robustness of this sex-biased microglial inflammatory program, we applied the same donor-level pseudobulk GSEA framework to two independent snRNA-seq cohorts, ROSMAP and the Netherlands Brain Bank (NBB). Both datasets validated the female-biased, disease-emergent enrichment of inflammatory pathways in microglia (**Figure 2E-F, Supplementary Figure 2B**). Thus, the microglial inflammatory response shift was reproducible across analytical frameworks and across independent human AD datasets.

Because IL6-JAK-STAT3 signaling emerged consistently within three independent cohorts of AD patients, we next asked whether *STAT3* expression is sex biased at the RNA level. Consistent with the pathway-level convergence, donor-level pseudobulk expression analysis showed that STAT3 RNA expression increased with disease progression in female astrocytes and microglia but not in males (Wilcoxon; astrocytes, p < 0.001; microglia, p < 0.01) (**Supplementary Figure 2C**). Thus, the female-biased neuroinflammatory program is accompanied by increased expression of the STAT3 transcriptional hub in key glial compartments. Having identified STAT3 as a reproducible human signal, we next asked whether this axis was conserved in an independent amyloid-driven model and whether it could be explained simply by sex differences in plaque exposure.

### 3.4 Female-biased STAT3 pathway activation is conserved across human AD and 5xFAD mice

To test whether STAT3-associated microglial inflammatory signature with AD is conserved, we examined STAT3 activation in the 5xFAD mouse model using spatial transcriptomics and immunofluorescence. Meta-analysis of a published Visium dataset from 5xFAD mice at 4, 6, 8 and 12 months of age showed that female 5xFAD mice had higher STAT3 target-module activity and higher DAM/microglial signature scores than age-matched male 5xFAD mice and wild-type controls (**Fig 3A, B**). STAT3 target activity remained relatively stable across age in wild-type mice but increased with age in 5xFAD mice, with the strongest increase observed in females (**Figure 3A,B**). This trajectory was accompanied by parallel increases in microglial and DAM scores, supporting the interpretation that the female-biased STAT3 signal reflects disease-progressive microglia inflammation (**Figure 3B).** Consistent with this relationship, STAT3 target-module activity strongly correlated with both plaque score and DAM/microglial score (R=0.89 and 0.8, respectively, **Figure 3C**). These correlations were significant in both males and females, suggesting that STAT3 signaling is associated with plaques and DAM microglia in general. To validate these spatial transcriptomic findings at the protein level, we performed immunofluorescence analyses of cortex and hippocampus from 5-month-old male and female 5xFAD mice. Consistent with the Visium analysis, females showed higher density of STAT3+ microglia (STAT3+IBA1+), a higher percentage of pSTAT3+ cells, and greater plaque density than males (**Figure 3D,E; Supplementary Figure 3A,B**).

**Figure 3.**
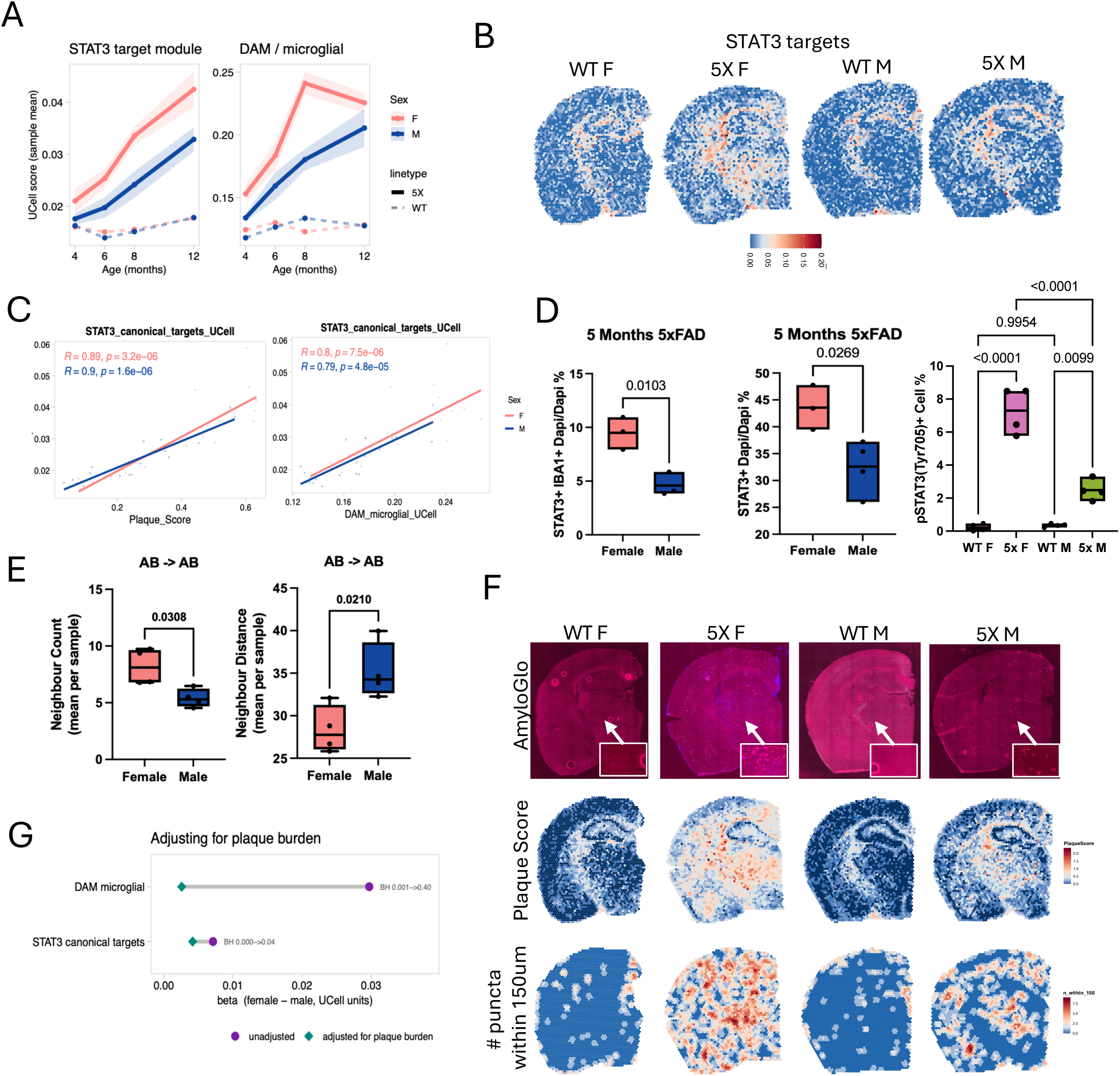
Cross species validation of STAT3 activation in AD microglia in 5xFAD mouse model. **A-C.** metanalyses of 5XFAD mouse Visium dataset (GSE233208). **A.** The trajectory of STAT3 targets module enrichment score across ages. Each point represents the mean and the ribbon represents +/− SEM., n = 5 mice per sex x genotype x age (5xFAD female 12 mo n = 7; WT female 12 mo n = 3). **B.** Representative spatial featureplots of STAT3 target enrichment scores in 8 months old mice. **C.** correlation between STAT3 target score and plaque score or DAM score. **G-E.** Immunofluorescence analyses of 5 months old 5X FAD female and male mice n=3 each showing **D.** % of pSTAT3, % of STAT3+ cells, or % of STAT3 and IBA1+ cells of all nuclei. P values from T tests or ANOVA with Tukey’s multiple comparisons test. **E**. Neighborhood analyses of amyloid beta plaques showing number of neighboring AB or the distance from the closest AB n= 4 mice each, p values represent Unpaired t test with Welch’s correction. **F.** Representative immunofluorescence and spatial featureplots from 8 Months old mice showing AmyloGlo staining, plaque module enrichment score and number of plaques within 150um diameter from the plaques. **G.** differences in ucell enrichment scores before and after correcting for plaque scores.

We next asked whether the female-biased STAT3/DAM signal could be explained by greater amyloid plaque exposure. Because the plaque-induced gene (PIG) signature overlaps with microglial and DAM markers, we leveraged the matched Amylo-Glo-stained Visium sections from the Miyoshi et al. dataset to quantify local plaque burden around each spot, including distance to the nearest plaque and the number of plaques within 100, 150, and 250 µm radii. This plaque-proximity analysis confirmed that female 5xFAD mice had higher local plaque burden than males (**Figure 3F, Supplementary Figure 3C**). Consistently, our independent amyloid-β immunofluorescence analysis at 5 months showed increased Aβ-to-Aβ neighborhood interactions in female 5xFAD mice, whereas male mice showed greater inter-plaque neighborhood distance (**Figure 3E**).

The transcriptional plaque-proximity score closely tracked image-based plaque measurements, correlating with both distance from plaques and the number of plaques within 150 µm in female and male 5xFAD mice (female, *R* = 0.77; male, *R* = 0.82), supporting its use as a spatial proxy for local plaque exposure (**Supplementary Figure 3D**). We therefore adjusted sex comparisons for plaque-proximity score to determine whether the female-biased inflammatory signals were explained by plaque burden. After adjustment, the DAM signature was no longer significantly different between females and males, indicating that the sex difference in the DAM-like response largely reflected greater plaque exposure in females. In contrast, the canonical STAT3 target module remained significantly female biased after plaque adjustment (**Figure 3G**). Thus, although female 5xFAD mice exhibit greater local plaque burden, plaque exposure alone does not explain the female-biased activation of the STAT3 pathway.

### 3.5 De novo microglial clustering reveals sex-divergent disease remodeling

Having established that STAT3 activation is female biased in human AD and remains female biased in 5xFAD mice after accounting for plaque proximity, we next asked which human microglial states carry this disease-associated program. This was particularly important because the broader DAM-like signal in the mouse spatial data was largely explained by plaque burden, whereas STAT3 remained sex biased after plaque adjustment. We therefore re-clustered the human microglial compartment de novo into 18 states (**Figure 4A**). Cluster identities were assigned using top differentially expressed markers, cross-validated against published microglial signatures from HuMiCA and ROSMAP, and Slingshot pseudotime analysis was performed to infer their differentiation trajectory (**Figure 4A-C; Supplementary Figure 4A,B; Supplementary Table 2**).

**Figure 4.**
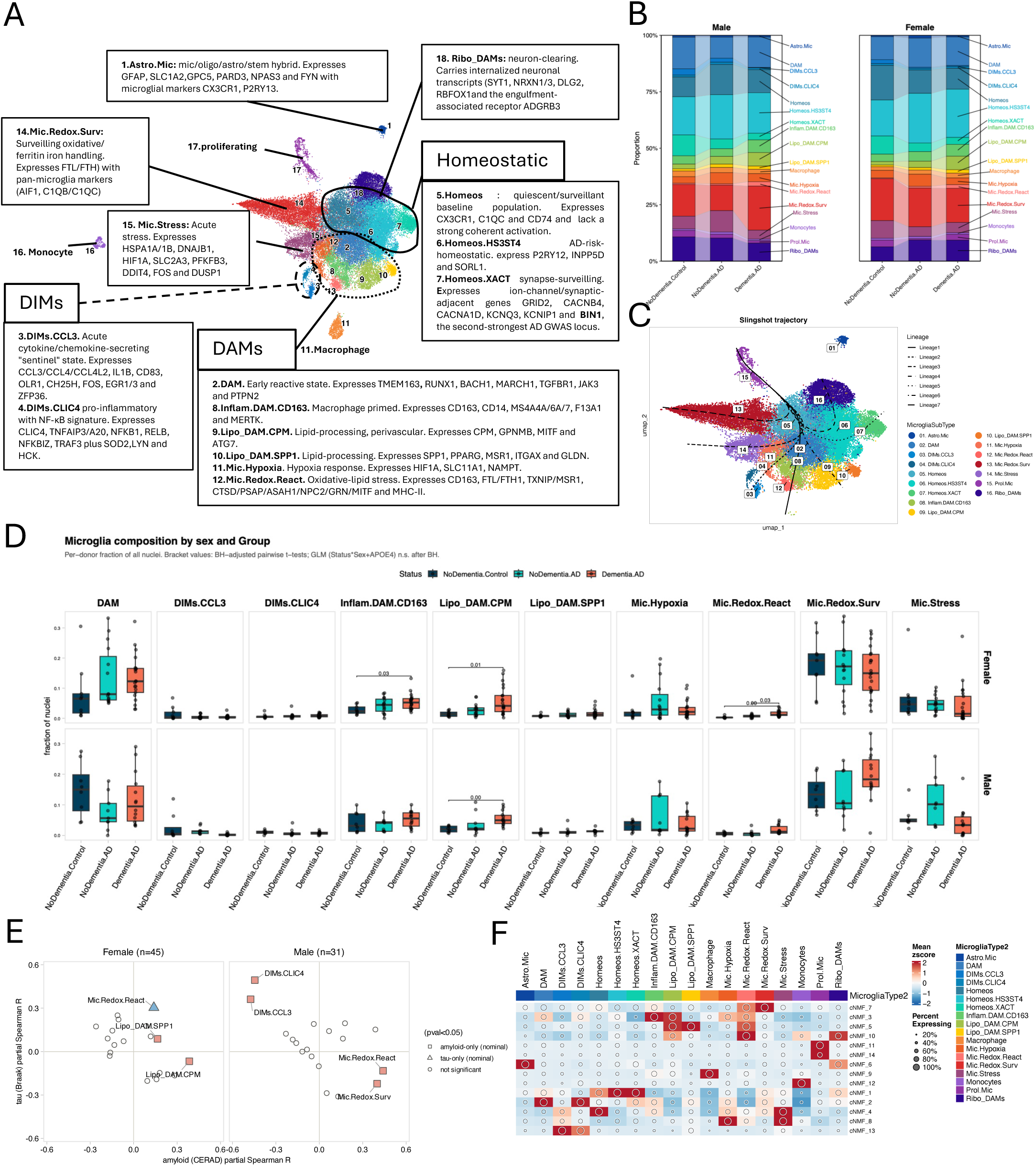
De novo microglial clustering resolves disease-emergent states and a larger homeostatic-to-myeloid shift in females. **A.** UMAP of the re-clustered microglial compartment, colored by annotated state identity (MicrogliaType). **B.** Stacked composition-trend plots of microglial state proportions across the Dementia.AD trajectory (NoDementia.Control, NoDementia.AD, Dementia.AD), shown separately for male (left) and female (right) donors. C. Slingshot trajectory for 7 lineages rooted at ‘Prol.Mic’, fit on the umap embedding. **D.** Per-donor microglia fractions by sex and cognitive status, modelled per cell type with a quasibinomial GLM adjusted for APOE and a sex by disease interaction. **E.** Partial Spearman correlation between donor-level composition (% of total microglia) and amyloid (CERAD score, x-axis) and tau (Braak stage, y-axis), faceted by sex. Each axis is residualized against the other pathology measure before correlation. Point shape indicates nominal (p<0.05) from a quasibinomial GLM (state fraction ∼ Braak + CERAD + APOE4). **F.** Heatmap of zscore-scaled mean usage score per cNMF. Dot size encodes the percent of cells expressing each module.

Unbiased pseudotime analysis organized microglia states along six trajectories. All trajectories passed through a homeostatic quiescent/surveillant hub state (Homeos; *CX3CR1, C1QC, CD74*). Among the homeostatic microglia, two additional populations were identified: an AD-risk-associated homeostatic state marked by *P2RY12, INPP5D, SORL1,* and *HS3ST4* (Homeos.HS3ST4), and a synapse-surveilling state enriched for neuronal and synaptic genes such as *GRID2, CACNB4, CACNA1D, KCNQ3, KCNIP1,* and *XACT* (Homeos.XACT). The pseudotime branched towards six DAM-like states: an early-reactive gateway state (DAM; *TMEM163, RUNX1, BACH1*) that served as a second pseudotime hub; a macrophage-primed CD163+ state (Inflam.DAM.CD163; *CD163, CD14, MS4A4A/6A/7, MERTK*); two lipid-processing states (Lipo_DAM.CPM; *CPM, GPNMB, MITF, and Lipo_DAM.SPP1; SPP1, PPARG, MSR1, ITGAX*); a hypoxia-response state (Mic.Hypoxia; *HIF1A, SLC11A1, NAMPT*); and a redox-reactive, oxidative-lipid-stress state (Mic.Redox.React; *CD163, FTL/FTH1, TXNIP, CTSD/PSAP/ASAH1/NPC2/GRN*). We also identified two disease-inflammatory microglial states: a cytokine/chemokine-producing state (DIMs.CCL3; *CCL3, CCL4, CCL4L2, IL1B, EGR1/2/*3) and an NF-κB-associated pro-inflammatory state (DIMs.CLIC4; *CLIC4, TNFAIP3, NFKB1, RELB*). Additional states included a surveilling redox and ferritin/iron-handling population (Mic.Redox.Surv; *FTL/FTH1, AIF1, C1QB/C1QC*), a multi-component acute-stress state (Mic.Stress; *HSPA1A/B/DNAJB1/HSPH1, HIF1A/SLC2A3/PFKFB3/DDIT4, FOS/DUSP1/ZFP36*), border-associated/perivascular macrophages (Macrophage; *MRC1, F13A1, LYVE1, CD200R1*), and infiltrating peripheral monocytes (Monocytes*; FCN1, VCAN*).

In addition to these canonical and disease-associated populations, we detected two noncanonical microglial states. The first, Astro.Mic, co-expressed microglial features together with astrocytic, oligodendrocyte-associated, and stem-cell-associated transcripts, including *GFAP, SLC1A2, GPC5, PARD3, and FYN*. The second, Ribo_DAM, named in relation to the Ribo-DAM1 population described in HuMiCA, was enriched for synaptic and neuronal transcripts, including *SYT1, NRXN1/3, DLG2, RBFOX1*, and *ADGRB3*, consistent with microglia containing internalized neuronal or synaptic material. To benchmark these annotations against an independent reference, we mapped our microglial states onto the pre-annotated ROSMAP microglia object using SCP::RunKNNMap, a K-nearest-neighbor-based reference mapping approach **(Supplementary Figure 4C)**. Most populations showed strong agreement with ROSMAP annotations, including Mic.Redox, monocytes, macrophages, and proliferating cells **(Supplementary Figure 4C).** In contrast, Astro.Mic and Ribo_DAM did not map to corresponding ROSMAP populations, likely because these cells were removed during prior filtering of the ROSMAP object. Multiple quality-control analyses supported retaining these clusters: both showed high ROGUE entropy-based homogeneity scores (>0.97), passed doublet filtering, and had feature and count distributions within the range observed for other microglial states **(Supplementary Figure 4D).**

### 3.6 Sex biased microglia states in AD progression

We next asked whether microglial state composition differed across sex and disease groups. Several disease-associated states expanded along the AD trajectory, but with distinct sex-specific patterns. Mic.Redox.React and Inflam.DAM.CD163 increased specifically in female donors between Control and Dementia.AD groups (Mic.Redox.React, padj = 0.0012; Inflam.DAM.CD163, padj = 0.026), whereas Lipo_DAM.CPM increased in both females and males (female, padj = 0.0068; male, padj = 0.0042) (**Figure 4D; Supplementary Figure 4E**). To determine whether these compositional shifts tracked neuropathologic burden, we performed partial Spearman correlations between microglial state abundance and Braak tau stage or CERAD amyloid score, after residualizing both variables against the remaining covariates. This analysis again highlighted Mic.Redox.React: in females, Mic.Redox.React was significantly associated with tau burden, whereas in males it was associated with amyloid burden(**Figure 4E**). Additional associations were also sex divergent, with DIM states negatively associated with amyloid in males and lipid-associated DAM states positively associated with amyloid in females (**Figure 4E**). Notably, STAT3 expression increased significantly in female but not male Dementia.AD donors within Mic.Redox.Surv, Ribo_DAM, and hypoxia-associated microglial populations compared with controls, linking female-biased STAT3 activation to multiple disease-associated microglial states (Supplementary Figure 4F).

To capture transcriptional programs that may cut across discrete cluster boundaries, we next applied consensus non-negative matrix factorization (cNMF), recovering 14 microglial gene programs (**Figure 4F; Supplementary Figure 4G,H**). These programs aligned closely with the de novo microglial states and included homeostatic(cNMF1), reactive TMEM163(cNMF2), primed CD163 (cNMF3), stress-associated regulatory (cNMF4), lipid-processing (cNMF5), astrocyte/glial-hybrid (cNMF6), ribosomal/translation (cNMF7), hypoxic/HIF1A metabolic (cNMF8), perivascular macrophage (cNMF9), oxidative phosphorylation/mitochondrial (cNMF10), cell-cycle (cNMF11&14), monocyte (cNMF12), and acute chemokine/cytokine modules (cNMF13). Several modules changed with disease in a sex-dependent manner. The CD163 module (cNMF3) and lipid-processing module (cNMF5) increased with disease in both sexes, whereas the oxidative phosphorylation/mitochondrial module (cNMF10) increased significantly in females but not males (female p = 0.005; male p = 0.08; **Supplementary Table 3**). Consistent with the cluster-level pathology associations, cNMF3 was positively correlated with amyloid burden in both sexes, while the chemokine/cytokine module cNMF13 was negatively associated with amyloid burden in males (**Supplementary Figure 4I**).

### 3.7 Sex-biased intercellular signaling links microglial remodeling to glial and vascular communication modules

Because de novo clustering suggested sex-divergent microglial remodeling, we next asked whether these state differences were accompanied by altered intercellular signaling. CellChat analysis across the six groups identified several signaling pathways with strong variation in total information flow (**Figure 5A**). TGFβ signaling was highest in female Dementia.AD and, together with GRN and ESAM, defined a female-leaning glial communication module that paralleled the female-biased inflammatory program we reported earlier (**Figure 5A; Supplementary Figure 5A,B**). We also observed divergent outgoing signaling from T/NK cells, with higher CD96, MHC-I, and PVR signaling in females with Dementia.AD and higher IL2 signaling in males with Dementia.AD (**Figure 5A; Supplementary Figure 5C**).

**Figure 5.**
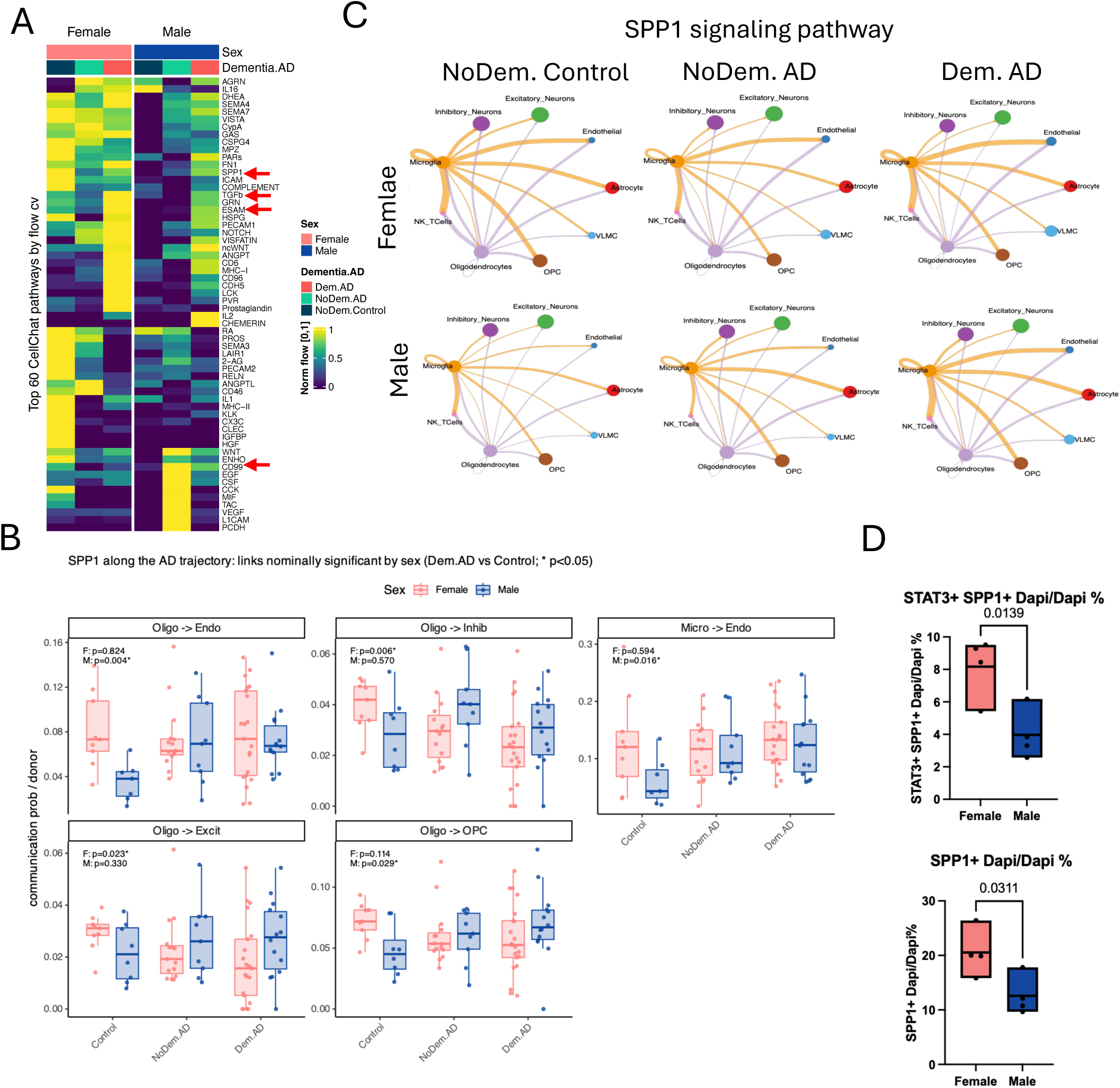
Sex-biased cell:cell signaling among microglia and other cell types in AD. **A.** Heatmap of the top 30 CellChat signaling pathways ranked by information-flow range across 6 conditions. Each value is the total communication probability summed across all sender-receiver pairs for that pathway, row-normalized to [0-1] so colour encodes which condition has relatively higher activity. Pathways are hierarchically clustered; conditions retain their input order. **B.** Circle plots of communication probability showing cross-condition comparison of the SPP1 signaling pathway across groups (p-value threshold: 0.05). **C.** Donor-level SPP1 communication probability. **D.** Immunofluorescence analyses of 5 months old 5X FAD female and male mice n=3 each showing SPP1+ or SPP1+,STAT3+ cells

CD99 showed the clearest male-biased communication pattern, consistent with previous reports^22^,. At the donor level, CD99 ligand-receptor links were the only communications significantly different between sexes in Dementia.AD after Benjamini-Hochberg correction, with endothelial and astrocyte senders driving the signal (**Figure 5A; Supplementary Figure 5D-E**). CD99 outgoing strength from these senders was higher in males and increased along the disease trajectory, becoming significant from NoDementia.AD onward (not significant in Control, p = 0.004 in NoDementia.AD, BH-adjusted p < 0.001 in Dementia.AD**, Supplementary Figure 5E**).

SPP1 signaling, a microglia-derived osteopontin pathway repeatedly implicated in AD^35–38^, showed a sex-divergent rather than simply female-biased pattern (**Figure 5A-C**). In females, SPP1 communication was high at baseline and declined with disease, whereas in males it was low at baseline and increased along the disease trajectory, leading to convergence between sexes in Dementia.AD (information flow: 1.31 in females versus 1.32 in males). Donor-level analysis further showed that the disease-associated increase in SPP1 signaling was male-specific and directed toward the vasculature, particularly through oligodendrocyte-and microglia-to-endothelial communication (nominal p < 0.05, **Figure 5B**), with no net sex difference in the dementia state.

Thus, at the global communication level, SPP1 appears to be a male-emergent vascular signal that increases with disease until it reaches the already high baseline observed in females. Since STAT3 is known to regulate SPP1 expression^39^, we asked STAT3+SPP1+ cells also show sex-biased abundance in 5xFAD mice. At 5 months, both the fraction of SPP1+ cells and the fraction of STAT3+SPP1+ double-positive cells were significantly higher in female than male 5xFAD mice (p < 0.0139; **Figure 5D**). This provides independent in situ support for a female-biased STAT3-SPP1 axis in an orthogonal amyloid-driven model.

We then asked whether a specific SPP1 vascular route was engaged in the human cohort. SPP1 ligand expression was concentrated predominantly in microglia, whereas the integrin receptor ITGA5 was enriched in endothelial cells and increased with disease (**Supplementary Figure 5F,G**). Decomposition of the SPP1 pathway into individual ligand-receptor pairs showed that the SPP1-ITGA5/ITGB integrin route was inferred only in dementia, while other SPP1 receptor routes were present across conditions. Thus, although total SPP1 communication is sex divergent and converges in Dementia.AD, its endothelial ITGA5/ITGB route is selectively enriched in the dementia state. Together, these findings link the STAT3-SPP1 program to a dementia-associated microglia-to-vasculature communication route, while distinguishing global sex-divergent SPP1 dynamics from the specific vascular mechanism active in dementia.

### 3.8 Cytokine-response profiling nominates divergent upstream drivers of female and male microglial programs

Finally, after defining the female-biased inflammatory programs, microglial state trajectories, and downstream communication architecture, we asked which upstream cytokine responses might drive these sex-divergent microglial programs. We performed in silico cytokine profiling using the human cytokine atlas^31^to infer which secreted cytokine induced sex-biased gene expression in AD microglia. Two cytokine-response signatures significantly separated females and males in Dementia.AD microglia (BH-adjusted p < 0.05; **Figure 6A**). The IL-17D response was female biased, increased along the disease trajectory, and peaked in female Dementia.AD microglia, whereas the IL-7 response was male biased and peaked in male Dementia.AD microglia (**Figure 6B**). Cross-dataset analysis in ROSMAP and NBB validated the same trend for IL-17D/IL-17-related enrichment (**Figure 6C**).

**Figure 6.**
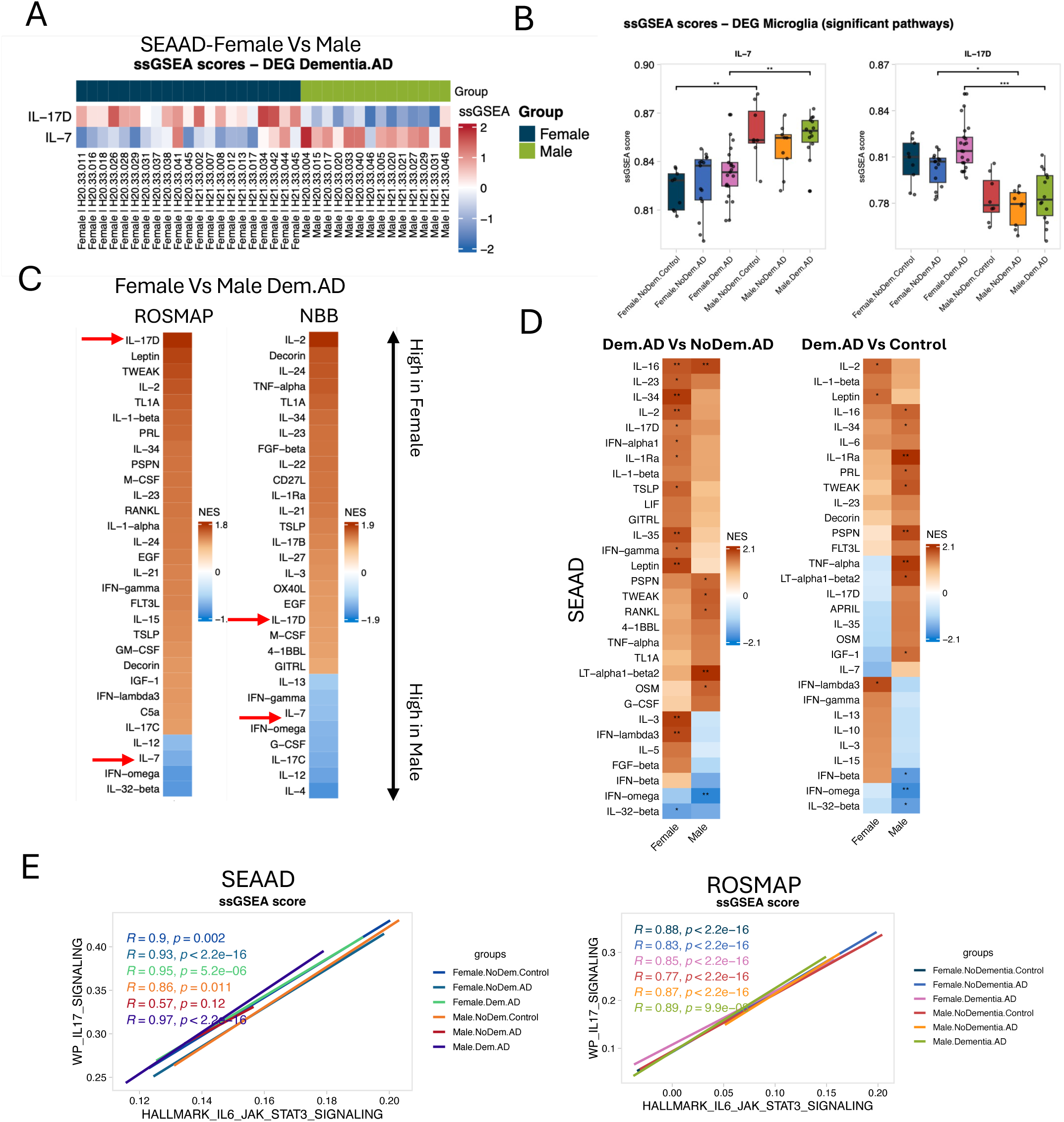
Human Cytokine Atlas analysis of DEG in donors with Dementia identifies sex-biased enrichment of IL17 signaling in females and IL7 signaling in males. **A.** Heatmap of single-sample GSEA (ssGSEA) enrichment scores in Dementia.AD cells for the Parse cytokine gene sets. Rows: the top significantly enriched signatures (BH-adjusted p < 0.05), IL-17D and IL-7; columns: individual donors, ordered by group (female, then male). Color: blue, low or negative enrichment; red, high or positive. Scores computed with GSVA::gsva() on log-CPM pseudobulk expression. **B.** Boxplots of ssGSEA scores for the IL-7 and IL-17D signatures in microglia across the six sex-by-stage groups; each point is one donor (raw p < 0.05). IL-7 peaks in male Dementia.AD; IL-17D peaks in female Dementia.AD. **C.** Heatmaps of donor-level pseudobulk GSEA Normalized Enrichment Scores for the cytokine signatures, for the Female vs Male contrasts, computed separately in ROSMAP and NBB datasets. Rows: top 30 signatures by max |NES|. **D.** Heatmaps of donor-level pseudobulk GSEA Normalized Enrichment Scores for the cytokine signatures, for the disease contrasts Dementia.AD vs NoDementia.AD (left) and Dementia.AD vs NoDementia.Control (right), computed separately in female and male donors (columns). Rows: top 30 signatures by max |NES|. Method: donor-level pseudobulk, edgeR TMM normalization, limma-voom moderated t-statistic pre-ranking, fgsea. NES > 0 = enriched in dementia. Stars: * BH < 0.05, ** < 0.01, *** < 0.001. **E.** Linear (lm) trend of BIOCARTA_IL17_PATHWAY vs HALLMARK_IL6_JAK_STAT3_SIGNALING ssGSEA scores, showing near perfect correlation between IL17 signaling and STAT3 signaling in microglia, colored by groups. SEAAD or ROSMAP Microglia cells were aggregated to one point per Donor.ID (mean of both axes) before fitting, so the trend reflects by donors rather than being weighted by cells per Donor.ID. Spearman’s rho and its p-value are shown.

Sex-stratified disease-transition analyses further supported a broad female-biased cytokine-response program (**Figure 6D**; **Supplementary Figure 6A**). In females, the Dementia.AD versus NoDementia.AD contrast was positively enriched for multiple inflammatory cytokine signatures, including IL-17D, IL-2, IL-34, IL-1β, IFN-α, and IFN-γ. In contrast, the corresponding male transition was enriched for TWEAK, RANKL, and OSM response signatures. In the Dementia.AD versus Control comparison, donor-level analyses identified significant IL-2, leptin, and IFN-λ enrichment in females, whereas males showed significant IL-16 and TNFα enrichment (Figure 6D; Supplementary Figure 6A). Direct expression analysis of cytokine, chemokine, and receptor genes, including IL, CCL/CCR, CXCL/CXCR, and CD-family genes, reinforced these group-specific patterns (**Supplementary Figure 6B**). Among all cytokine-response signatures tested, IL-2 was conserved across SEA-AD, ROSMAP, and NBB in both female and male Dementia.AD versus Control comparisons (**Figure 6D; Supplementary Figure 6C**).

Together, these analyses identify IL-17D/IL-17-related responsiveness as a reproducible female-biased cytokine-response program that emerges with AD dementia. This finding is notable because IL-17 signaling has been implicated in AD pathology in both human and mouse studies and can engage STAT3-dependent inflammatory programs. Consistent with this link, IL-17-related and STAT3 pathway scores were strongly correlated in AD microglia in SEA-AD and validated in ROSMAP, particularly in females (R > 0.9; **Figure 6E**). These results support a close association between IL-17-related cytokine responsiveness and the female-biased STAT3 activation signature identified here.

## 4. DISCUSSION

In this study, we used sex-stratified single-nucleus, spatial transcriptomic, cytokine-response, and protein-level analyses to define how biological sex is associated with the inflammatory response due to AD progression from without dementia to AD-associated dementia. The central finding is that female donors with Dementia.AD exhibit a coordinated, disease-emergent neuroinflammatory program spanning microglia, astrocytes, OPCs, and vascular-associated compartments. Unexpectedly, the microglial inflammatory signal did not simply appear in females with disease but reversed polarity along the trajectory. In cognitively normal donors without AD pathology (Control), the entire neuroinflammatory block was significantly higher in males than females (all six programs BH < 0.05). This male bias attenuated in NoDementia.AD, where IL6-JAK-STAT3 and complement no longer differed by sex while the interferon and TNFα responses remained male-biased, and then inverted to a strong female bias in Dementia.AD (**Figure 2A, Supplementary Figure 2A**). This disease-stage-dependent reversal suggests that sex differences in AD neuroinflammation are dynamic and context dependent, rather than fixed differences in immune tone. It also supports a model in which the transition from neuropathology without dementia to clinical dementia is accompanied by a coordinated elevation of inflammation in females but not in males.

A major implication of these findings is that STAT3 may represent a highly reproducible female-biased inflammatory hub across datasets and species. STAT3 expression itself increased with disease in female microglia and astrocytes but not in males, consistent with a cell-type-spanning activation axis rather than a microglia-only phenomenon. This interpretation is supported by prior work showing that AD neuroinflammation is shaped by sex-biased microglial responses and by broader sex-dependent mechanisms involving inflammation, metabolism, sex hormones, and chromosomal biology. It is also consistent with emerging views of STAT3 as a central, cell-type-dependent mediator of AD-associated neuroinflammatory and neurodegenerative processes.

Further, our mouse model (5xFAD) analyses extend this inference beyond the human cohort and help separate STAT3 activation from amyloid plaque burden. 5-month-old female 5xFAD mice showed higher STAT3 signal than males, and spatial transcriptomic reanalysis demonstrated an age-dependent increase in STAT3 target activity that was most pronounced in female 5xFAD sections. Although female mice also showed greater plaque burden, the plaque-adjusted analysis was informative: DAM scores were no longer significantly sex biased after adjustment for plaque proximity, whereas canonical STAT3 target activity remained significantly female biased. This distinction suggests that part of the broader DAM-like response reflects local plaque exposure, while the STAT3 axis captures an additional sex-biased inflammatory component not explained by plaque burden alone.

De novo microglial clustering refined this model by showing that the female-biased STAT3 program is embedded within separate subsets of microglia. Several disease-associated states expanded along the AD trajectory, but the strongest sex-specific changes occurred in redox-reactive and CD163+ macrophage-primed microglia, which increased preferentially in female donors. Moreover, partial pearson correlation revealed that while redox-reactive state is significantly associated with amyloid burden in males, it’s amyloid independent in females. More importantly, redox-reactive cells were positively associated with tau pathology only in females.

Cell-cell communication analyses placed these microglial changes within a broader glial-vascular signaling architecture. Consistent with prior work, CD99 emerged as the most robust male-biased donor-level communication axis in Dementia.AD, driven primarily by endothelial and astrocyte senders. This pattern is consistent with the pseudoautosomal location of CD99 and suggests that dosage-related signaling may become more apparent during disease progression. In contrast, TGFβ, GRN, and ESAM defined a female-leaning glial communication module that paralleled the female-biased inflammatory program. We also observed sex-divergent signaling involving the T/NK-cell compartment, with CD96, MHC-I, and PVR enriched in female Dementia.AD and IL2 enriched in male Dementia.AD. Notably, upstream cytokine-response profiling identified IL-2 as a conserved dementia-associated signature across SEA-AD, ROSMAP, and NBB in both sexes, suggesting that some cytokine-response programs accompany AD dementia broadly, even when individual communication routes are sex biased.

The cytokine-response analysis provides a candidate upstream explanation for the sex-divergent inflammatory routes observed in microglia. Female Dementia.AD microglia were enriched for IL-17D and interferon-linked response programs, whereas male Dementia.AD microglia showed stronger enrichment of an IL-7-associated response. This dichotomy is mechanistically consistent with the neuroinflammatory architecture identified above: IL-17 signaling can converge on NF-κB and STAT3, the female-biased inflammatory hub, whereas IL-7 signals primarily through JAK-STAT5. Together, these analyses nominate distinct upstream cytokine programs associated with sex-divergent microglial remodeling, including a female-biased IL-17/interferon-associated program linked to STAT3-centered inflammation and a separate male-biased IL-7-associated axis. The female IL-17D signal is particularly notable because it connects the dementia-associated microglial program to IL-17 receptor signaling through IL17RA/B/C, positioning IL-17-related signaling as a candidate driver of the female-biased response.

This interpretation is supported by prior evidence implicating IL-17 signaling in AD and related neuroinflammatory contexts^40,41^. In mouse models, IL-17 has been linked to cognitive decline, synaptic deficits, and neuroinflammation, and neutralization of IL-17 can reduce amyloid-induced inflammatory and memory phenotypes. Notably, Brigas et al. ^40^reported that meningeal γδT cells produce IL-17 in the 3xTg-AD model and that IL-17 expression was higher in females than males, consistent with the sex-biased IL-17-related signal observed here. Human studies also support a broader IL-17/Th17 axis in AD, including elevated peripheral IL-17-family cytokines, increased Th17 differentiation markers, and higher circulating Th17-cell proportions or IL-17/IL-23 levels in patients with AD^42,43^. Together with studies linking Th17 and γδT-cell biology to meningeal inflammation and blood-brain barrier integrity, these findings suggest that IL-17-related signaling may connect peripheral or border-associated immune programs to glial inflammatory states in AD. However, the cell types that produce and respond to IL-17 in the AD brain, and whether these interactions are sex biased, remain unresolved and require direct experimental testing.

Several limitations should be considered. First, the human analyses are based on postmortem tissue and therefore capture associations among sex, disease stage, and transcriptional state rather than causal mechanisms. Second, although donor-level pseudobulk analysis reduces pseudoreplication and the models accounted for age and APOE genotype, residual confounding by unmeasured clinical, hormonal, vascular, medication, or agonal factors cannot be excluded. Third, because the human analyses focused primarily on DLPFC, the extent to which these sex-biased programs generalize to other vulnerable brain regions remains to be determined. Fourth, the cytokine atlas was generated from peripheral CD14 monocytes rather than human microglia, so cytokine-response enrichment should be viewed as hypothesis-generating. Fifth, 5xFAD provides an amyloid-driven model for cross-species validation but does not capture the full spectrum of human AD, including tau pathology, cognitive heterogeneity, and aging-related comorbidities. Finally, CellChat and ligand-receptor analyses infer communication potential from transcript abundance and require functional validation.

## 5. CONCLUSION

Overall, our findings support a model in which biological sex shapes not only the magnitude of neuroinflammation in AD, but also the inflammatory route through which neuropathology progresses to dementia. In females, AD dementia is associated with a STAT3-centered program linking IL-17/interferon cytokine responsiveness, redox-reactive and CD163+ microglial remodeling, and glial-vascular communication. In males, the disease-associated inflammatory architecture is comparatively less STAT3-centered and includes distinct CD99 and IL-7-linked signaling features. This may help explain why sex differences in AD neuroinflammation vary across cohorts, disease stages, and model systems. It also suggests that immunomodulatory or glial-targeted therapies may need to account for sex, disease stage, and the specific inflammatory circuit engaged.

## Acknowledgements

This work was supported by generous gifts from Mr. Williams, Carolyn and Platt Davis, the MC Linn Family Foundation Fund, and the Houston Methodist Chair in Neurodegenerative Research Endowment fund (KY).

