## Supplementary Figures for "Female-Biased IL-17–STAT3 Signaling Marks Dementia-Associated Microglial Inflammation in Alzheimer’s Disease"

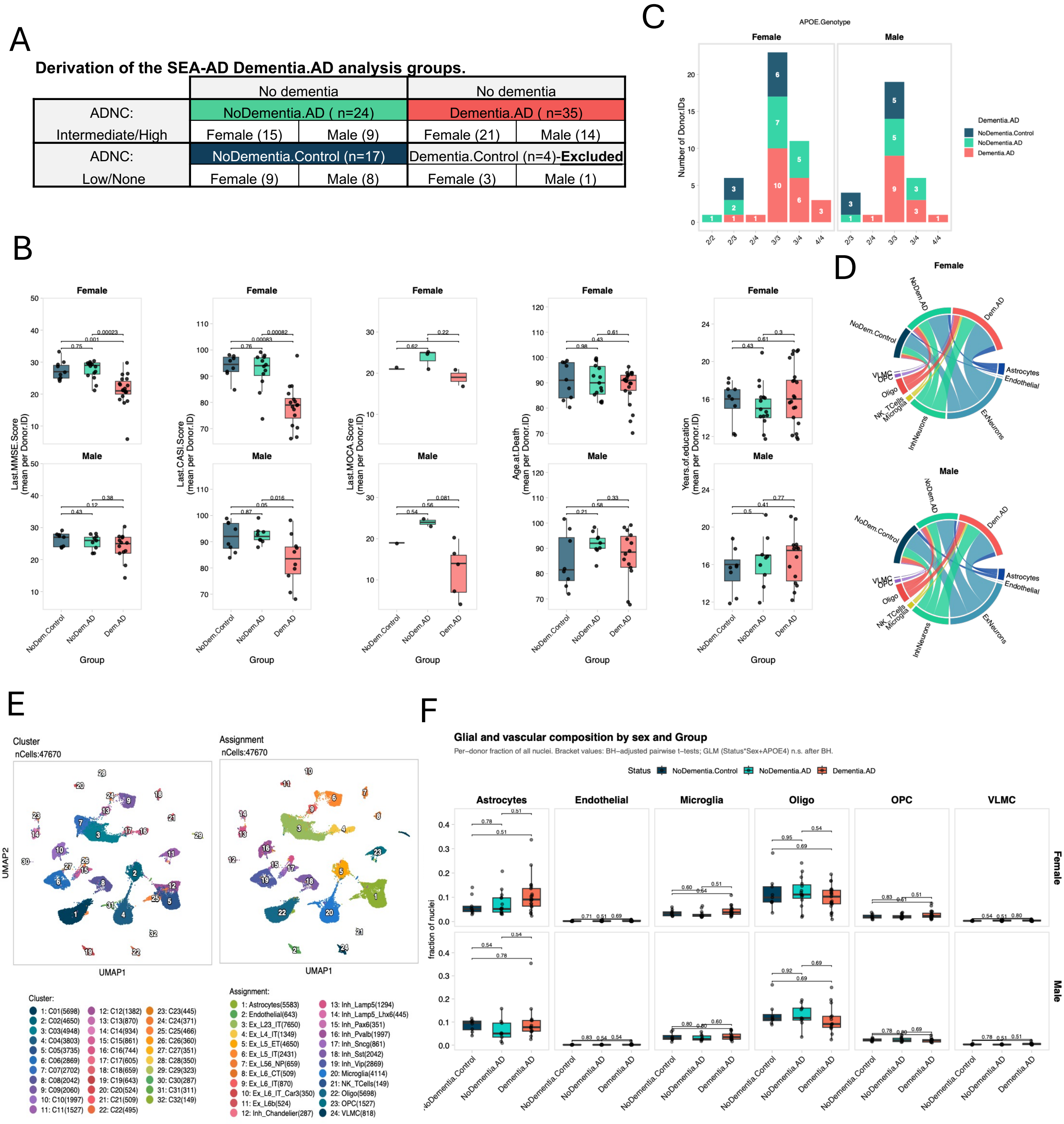
A
Derivation of the SEA-AD Dementia.AD analysis groups.
No dementia
No dementia
ADNC: Intermediate/High
NoDementia.AD ( n=24)
Dementia.AD ( n=35)
Female (15)
Male (9)
Female (21)
Male (14)
ADNC: Low/None
NoDementia.Control (n=17)
Dementia.Control (n=4)-Excluded
Female (9)
Male (8)
Female (3)
Male (1)
B
Female
Male
Last.MMSE.Score (mean per Donor.ID)
Last.CASI.Score (mean per Donor.ID)
Last.MOCA.Score (mean per Donor.ID)
Age.at.Death (mean per Donor.ID)
Years.of.education (mean per Donor.ID)
NoDem.Control
NoDem.AD
Dem.AD
Group
C
APOE.Genotype
Female
Male
Number of Donor.IDs
Dementia.AD
NoDementia.Control
NoDementia.AD
Dementia.AD
D
Female
Male
NoDem.AD
Dem.AD
NoDem.Control
Astrocytes
Endothelial
VLMC
OPC
Oligo
NK_TCells
Microglia
InhNeurons
ExNeurons
E
Cluster
nCells:47670
Assignment
nCells:47670
UMAP1
UMAP2
Cluster:
1: C01(5698)
2: C02(4650)
3: C03(4948)
4: C04(3803)
5: C05(3735)
6: C06(2869)
7: C07(2702)
8: C08(2042)
9: C09(2060)
10: C10(1997)
11: C11(1527)
12: C12(1382)
13: C13(870)
14: C14(934)
15: C15(861)
16: C16(744)
17: C17(605)
18: C18(659)
19: C19(643)
20: C20(524)
21: C21(509)
22: C22(495)
23: C23(445)
24: C24(371)
25: C25(466)
26: C26(360)
27: C27(351)
28: C28(350)
29: C29(323)
30: C30(287)
31: C31(311)
32: C32(149)
Assignment:
1: Astrocytes(5583)
2: Endothelial(643)
3: Ex_L23_IT(7650)
4: Ex_L4_IT(1349)
5: Ex_L5_ET(4650)
6: Ex_L5_IT(2431)
7: Ex_L56_NP(659)
8: Ex_L6_CT(509)
9: Ex_L6_IT(870)
10: Ex_L6_IT_Car3(350)
11: Ex_L6b(524)
12: Inh_Chandelier(287)
13: Inh_Lamp5(1294)
14: Inh_Lamp5_Lhx6(445)
15: Inh_Pax6(351)
16: Inh_Pvalb(1997)
17: Inh_Sncg(861)
18: Inh_Sst(2042)
19: Inh_Vip(2869)
20: Microglia(4114)
21: NK_TCells(149)
22: Oligo(5698)
23: OPC(1527)
24: VLMC(818)
F
Glial and vascular composition by sex and Group
Per−donor fraction of all nuclei. Bracket values: BH−adjusted pairwise t−tests; GLM (Status*Sex+APOE4) n.s. after BH.
Status
NoDementia.Control
NoDementia.AD
Dementia.AD
Astrocytes
Endothelial
Microglia
Oligo
OPC
VLMC
Female
Male
fraction of nuclei
NoDementia.Control
NoDementia.AD
Dementia.AD

G

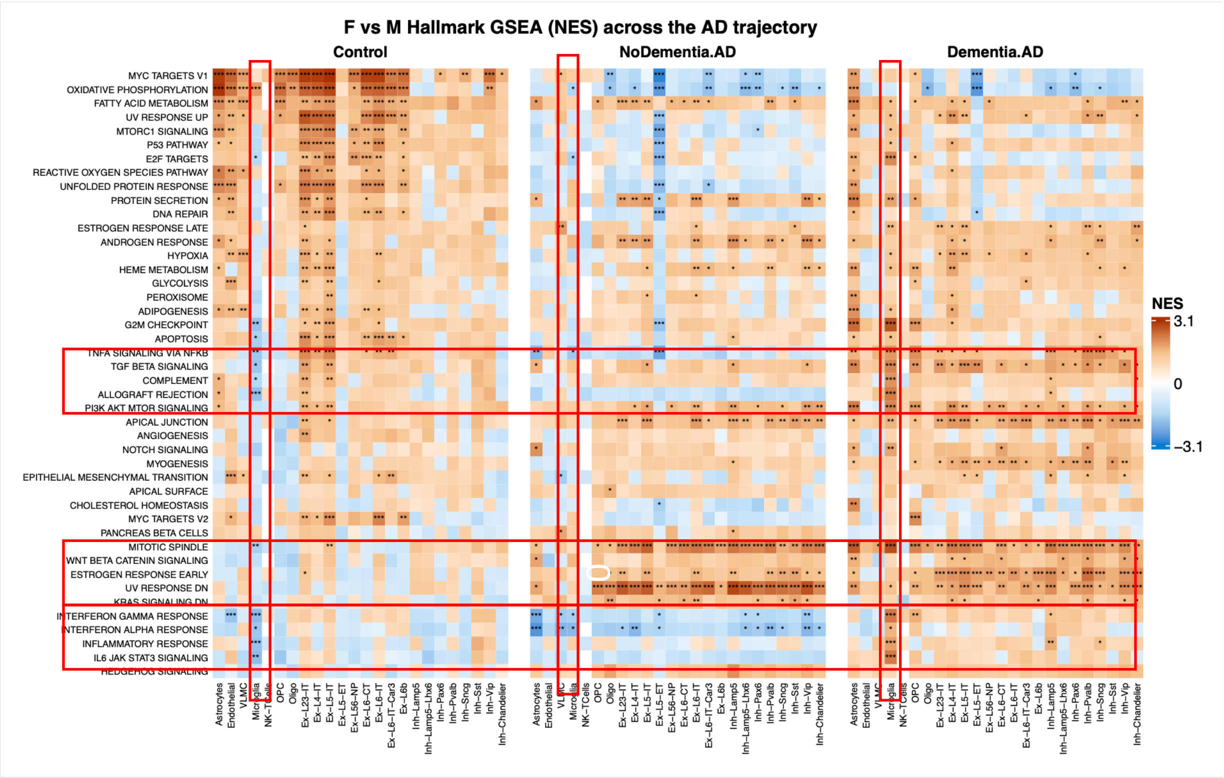


**Supplementary Figure 1: A. Derivation of the SEA-AD analysis groups.** Donors were classified by crossing two independent criteria: clinical cognitive status (No dementia vs. Dementia) and AD neuropathologic change (ADNC), the latter derived from Thal amyloid phase, Braak neurofibrillary tangle stage, and CERAD neuritic plaque score via the NIA-AA "ABC" consensus criteria, collapsed to Low/None vs. Intermediate/High. Crossing these axes yields four groups: NoDementia.Control (no dementia, no/low AD pathology; n=17, 9 female/8 male), NoDementia.AD (no dementia despite AD pathology, a resilient state; n=24, 15 female/9 male), Dementia.AD (dementia with AD pathology; n=35, 21 female/14 male), and a fourth cell (dementia without AD pathology, i.e. non-AD dementia; n=4, 3 female/1 male) that is excluded from the analysis cohort so that Dementia.AD reflects only clinically-and-pathologically confirmed Alzheimer's disease. Three additional reference brains (no cognitive or neuropathologic data collected) are excluded upstream of this classification. The three retained groups total 76 donors (45 female, 31 male). **B.** Summary of Last.MMSE.Score, Last.CASI.Score, Last.MOCA.Score, Age.at.Death, Years.of.education from 76 donors. Each dot = mean log-normalized expression for one Donor.ID, grouped by Group, rows by Sex. Box: median, IQR; whiskers: 1.5x IQR. Statistical comparisons: Wilcoxon rank-sum test on donor-level means. **C.** Stacked bar chart of summarizing the distribution of APOE.Genotype (deduplicated to one row per Donor.ID). Y-axis shows Number of Donor.IDs. Bars are stacked and colored by Dementia.AD, with each segment labeled with its count. Rows are split by Sex. **D.** Chord diagram showing the flow of cells across GlobalCellTypes -> Group. Arc width is proportional to cell count. Separate diagrams arranged in a 1-column grid, one per level of Sex. **E.** Single-nucleus landscape of the SEA-AD cohort. UMAP of 1,598,085 nuclei from 76 donors, integrated with Harmony across donor identity,library prep method; Louvain clustering (resolution 0.5) resolved 32 clusters annotated to 24 cell types. **F.** Per-donor glial and vascular fractions by sex and cognitive status, modelled per cell type with a quasibinomial GLM adjusted for APOE and a sex by disease interaction, and confirmed by Dirichlet regression. Microglial and astrocytic fractions trend higher in dementia, but no fraction differs significantly by sex, disease or APOE after BH correction. **G.** Heatmap of pseudobulk (Σ raw counts per donor × cell type → DESeq2 with covariates APOE, age, PMI) GSEA Normalized Enrichment Scores (Hallmark) for contrast F_vs_M in Control, NoDementia.AD or Dementia.AD across cell types. Rows = union of top 30 gene sets by max |NES| across cell types. Method: donor-level pseudobulk, edgeR TMM normalization, limma-voom moderated t-statistic pre-ranking, fgsea. NES > 0 = enriched in the first term of the contrast. Overlaid significance stars: * BH < 0.05, ** < 0.01, *** < 0.001.B. Summary NES heatmap for Microglia cells (Hallmark database). Rows = union of top 30 pathways per contrast, filtered to max |NES| >= 1. Columns = all contrasts tested. Stars: * BH < 0.05, ** < 0.01, *** < 0.001.

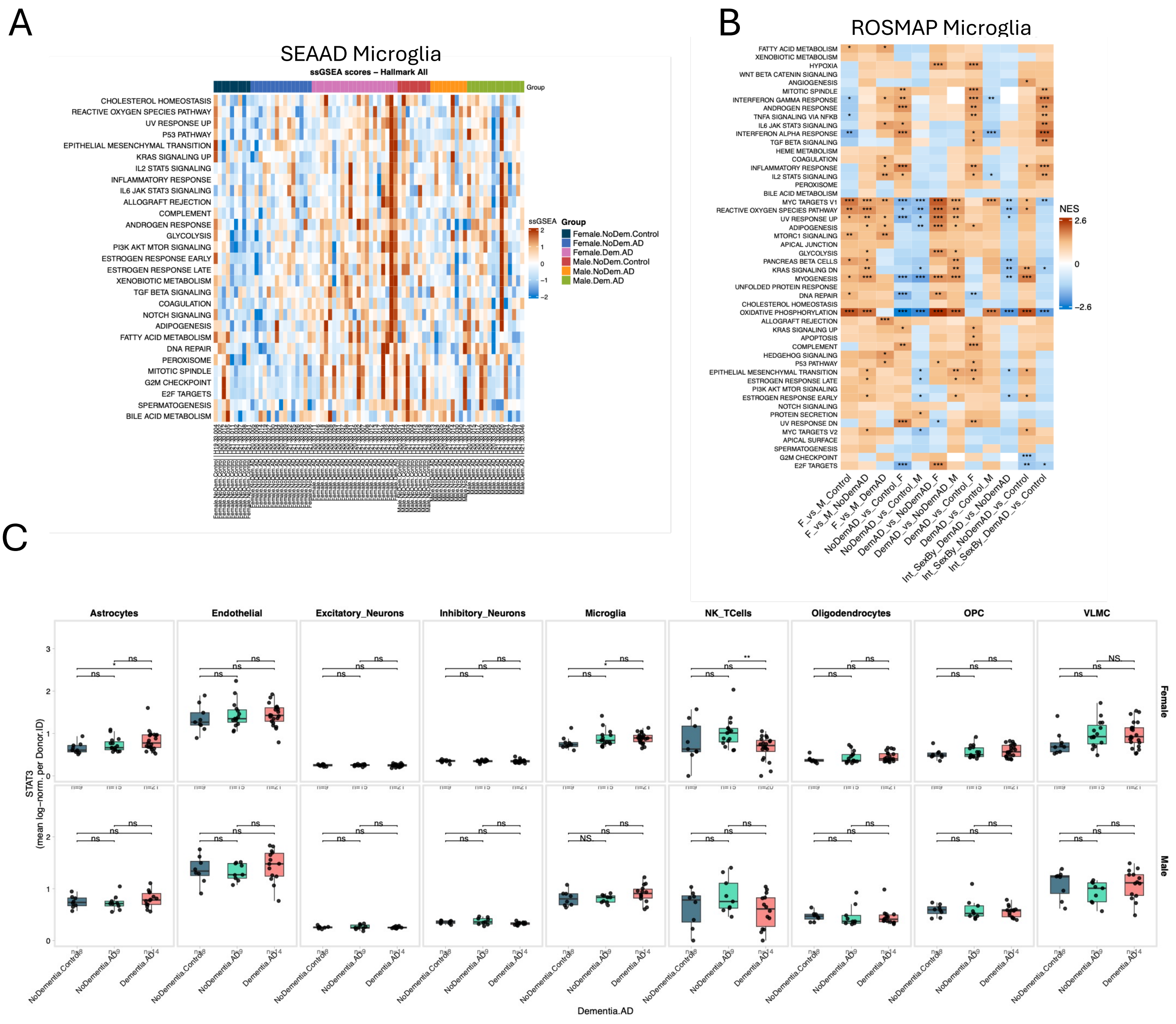


**Supplementary Figure 2. A female-biased, disease-emergent neuroinflammatory program across glia and vasculature. A.** Heatmap of single-sample GSEA (ssGSEA) enrichment scores for top significantly differentially enriched pathways in SEAAD Microglia cells (Hallmark database; BH-adjusted p < 0.05). Each column representa the means across individual donors. **B.** Pseudobulk GSEA NES heatmap (Hallmark) in ROSMAP Microglia, contrast: Int_SexBy_DemAD_vs_Control. Dataset: 75,265 cells. Rows = top 30 gene sets by max |NES| across cell types (columns). Method: edgeR TMM normalization, limma-voom moderated t-statistic pre-ranking, fgsea. Positive NES = enriched in the first term of the contrast. Stars: * BH < 0.05, ** < 0.01, *** < 0.001. **C.** Pseudobulk box plot of STAT3. Each dot represents the mean log-normalized expression for one Donor.ID (n = 76 donors), grouped by Dementia.AD, split by global cell type, rows by Sex. Box plot elements: centre line = median; box limits = 25th-75th percentile (IQR); whiskers extend to the furthest observation within 1.5x IQR from the box; outliers beyond this range are not shown. Statistical comparisons: Wilcoxon rank-sum test on donor-level means.

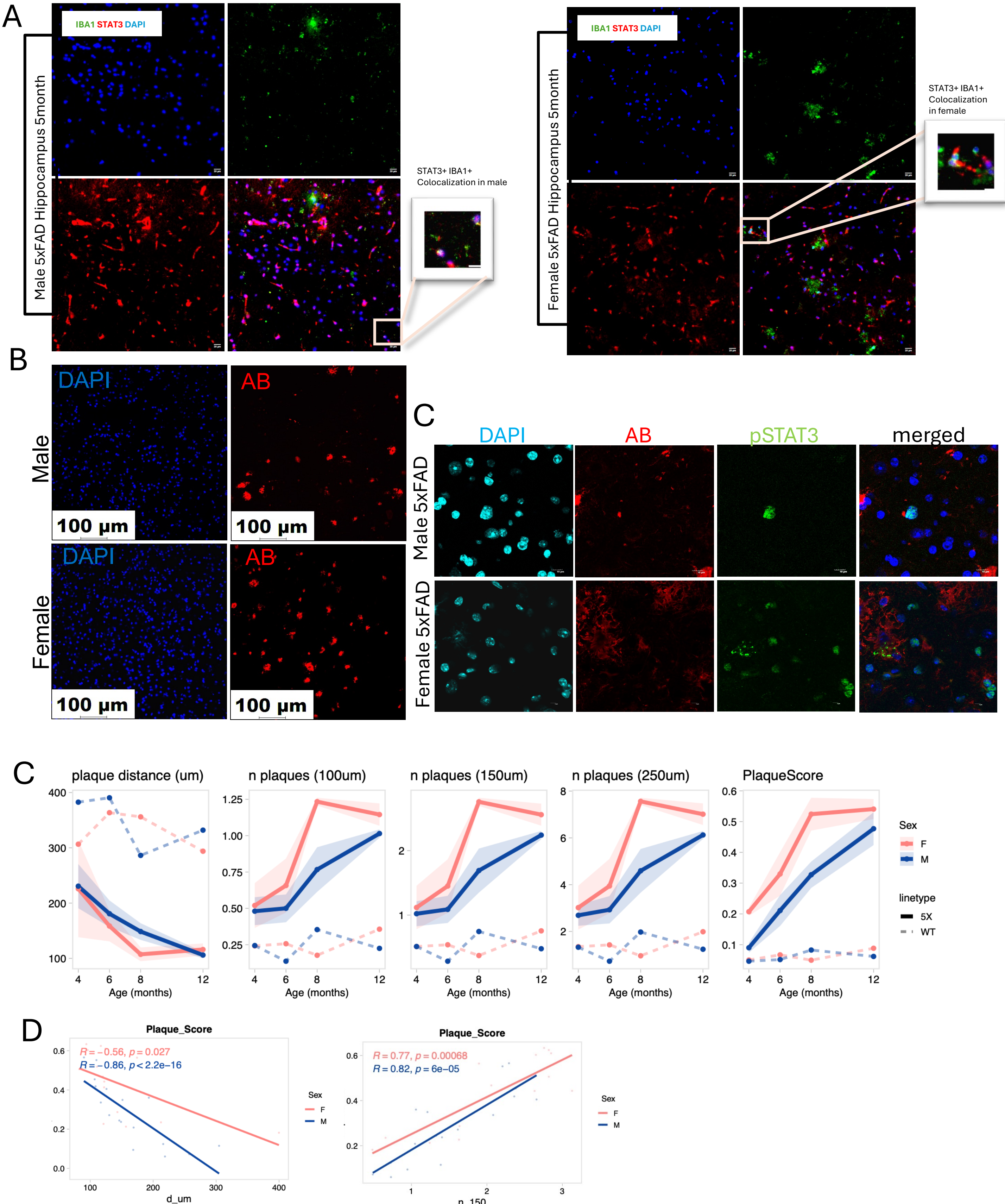


**Supplementary Figure 3. Multi-omics analyses of 5xFAD mouse model. A.** representative immunofluorescence images of STAT3 and IBA1 staining of 5 months old 5XFAD mice. **B.** representative immunofluorescence images of AB staining of 5 months old 5XFAD mice. **C-D.** metanalyses of 5XFAD mouse Visium dataset (GSE233208). **C** Line graphs showing the trajectory of distance of counted plaques, number of plaques within 100, 150 and 250 um diameter and the plaque module enrichment score across ages. Each point represents the mean and the ribbon represents +/- SEM., (4mo n= 4 WT each, 3 5XFAD each, 6mo n= 4 WT each, 3 5XFAD F and 5 M, 8mo 3 F each, 5 M each, 12mo 4 M each and 7 F 5XFAD, 3 F WT). **D.** correlation between transcriptional plaque score and distance from image based detected plaques.

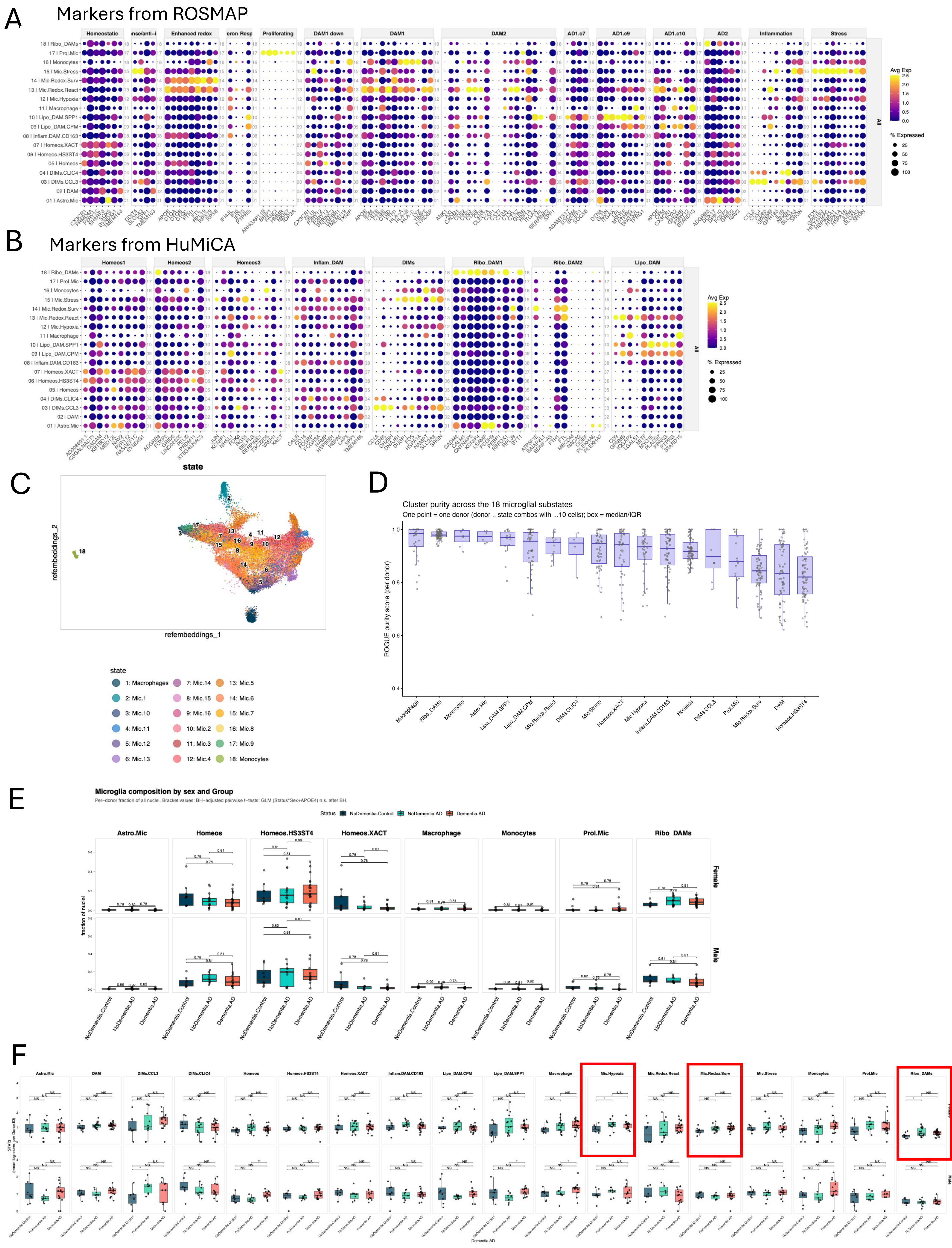
A
Markers from ROSMAP
Homeostatic
Enhanced redox
Proliferating
DAM1 down
DAM1
DAM2
AD1.c7
AD1.c9
AD1.c10
AD2
Inflammation
Stress
Avg Exp
% Expressed
18 | Ribo_DAMs
17 | Prol.Mic
16 | Monocytes
15 | Mic.Stress
14 | Mic.Redox.Surv
13 | Mic.Redox.React
12 | Mic.Hypoxia
11 | Macrophage
10 | Lipo_DAM.SPP1
09 | Lipo_DAM.CPM
08 | Inflam.DAM.CD163
07 | Homeos.XACT
06 | Homeos.HS3ST4
05 | Homeos
04 | DIMs.CLIC4
03 | DIMs.CCL3
02 | DAM
01 | Astro.Mic
B
Markers from HuMiCA
Homeos1
Homeos2
Homeos3
Inflam_DAM
DIMs
Ribo_DAM1
Ribo_DAM2
Lipo_DAM
Avg Exp
% Expressed
C
state
refembeddings_2
refembeddings_1
1: Macrophages
2: Mic.1
3: Mic.10
4: Mic.11
5: Mic.12
6: Mic.13
7: Mic.14
8: Mic.15
9: Mic.16
10: Mic.2
11: Mic.3
12: Mic.4
13: Mic.5
14: Mic.6
15: Mic.7
16: Mic.8
17: Mic.9
18: Monocytes
D
Cluster purity across the 18 microglial substates
One point = one donor (donor .. state combos with ...10 cells); box = median/IQR
ROGUE purity score (per donor)
Macrophage
Ribo_DAMs
Monocytes
Astro.Mic
Lipo_DAM.SPP1
Lipo_DAM.CPM
Mic.Redox.React
DIMs.CLIC4
Mic.Stress
Homeos.XACT
Mic.Hypoxia
Inflam.DAM.CD163
Homeos
DIMs.CCL3
Prol.Mic
Mic.Redox.Surv
DAM
Homeos.HS3ST4
E
Microglia composition by sex and Group
Per–donor fraction of all nuclei. Bracket values: BH–adjusted pairwise t–tests; GLM (Status*Sex+APOE4) n.s. after BH.
Status
NoDementia.Control
NoDementia.AD
Dementia.AD
Astro.Mic
Homeos
Homeos.HS3ST4
Homeos.XACT
Macrophage
Monocytes
Prol.Mic
Ribo_DAMs
Female
Male
fraction of nuclei
F
Astro.Mic
DAM
DIMs.CCL3
DIMs.CLIC4
Homeos
Homeos.HS3ST4
Homeos.XACT
Inflam.DAM.CD163
Lipo_DAM.CPM
Lipo_DAM.SPP1
Macrophage
Mic.Hypoxia
Mic.Redox.React
Mic.Redox.Surv
Mic.Stress
Monocytes
Prol.Mic
Ribo_DAMs
Dementia.AD

G

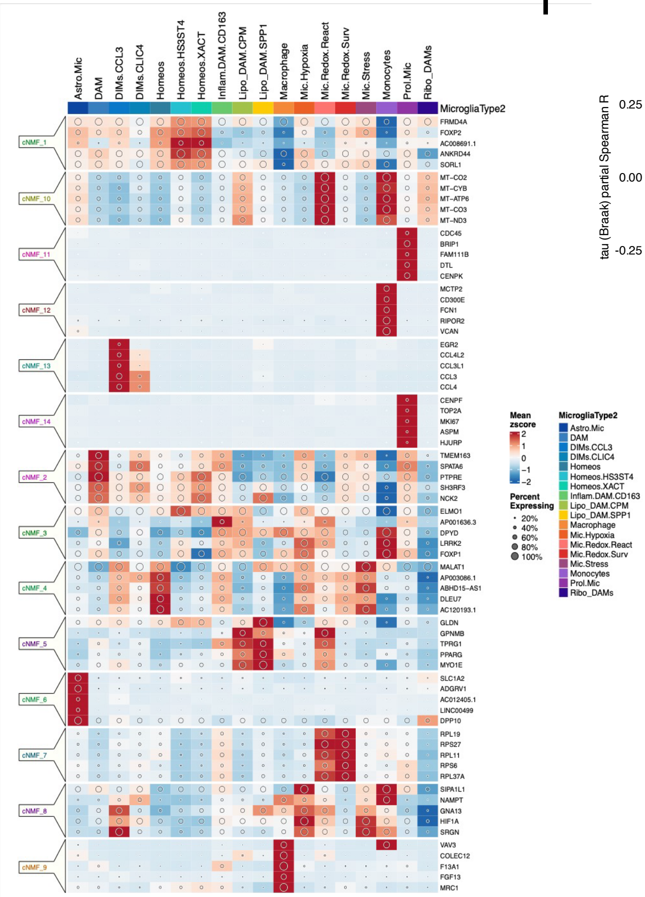

I

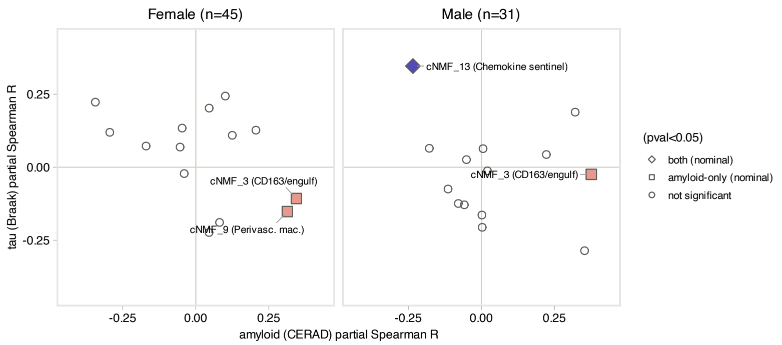


H

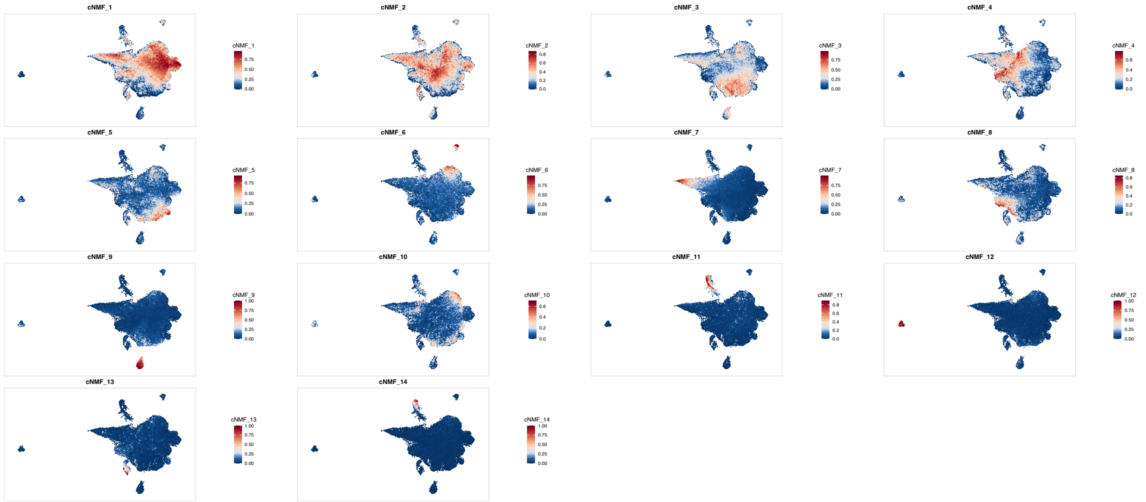

**Supplementary Figure 4**. **De novo microglial clustering. A.** Dot plot of Microglia markers from the ROSMAP manuscript. **B.** Dot plot of Microglia markers from the HuMiCa manuscript. Dot size denotes the percent of cells expressing each gene; dot color denotes scaled mean expression. **C.** K-nearest-neighbor-based reference mapping of ROSMAP microglia object using SEAAD embedding. **D.** Per donor ROGUE entropy-based homogeneity scores. **E.** Per-donor microglia fractions by sex and cognitive status, modelled per cell type with a quasibinomial GLM adjusted for APOE and a sex by disease interaction, and confirmed by Dirichlet regression. **F.** Pseudobulk box plot of STAT3. Each dot represents the mean log-normalized expression for one Donor.ID (n = 76 donors), grouped by Dementia.AD, split by global cell type, rows by Sex. Box plot elements: centre line = median; box limits = 25th-75th percentile (IQR); whiskers extend to the furthest observation within 1.5x IQR from the box; outliers beyond this range are not shown. Statistical comparisons: Wilcoxon rank-sum test on donor-level means. **G-I.** cNMF analysis of microglia modules, **G.** Heatmap of zscore-scaled mean expression for top 5 genes per cNMF. Dot size encodes the percent of cells expressing each gene. **H.** **F**eatureplots showing enrichment of each module. **I.** Partial Spearman correlation between donor-level cNMF usage score and amyloid (CERAD score, x-axis) and tau (Braak stage, y-axis), faceted by sex. Each axis is residualized against the other pathology measure before correlation. Point shape indicates nominal (p<0.05) from a quasibinomial GLM (state fraction ~ Braak + CERAD + APOE4).

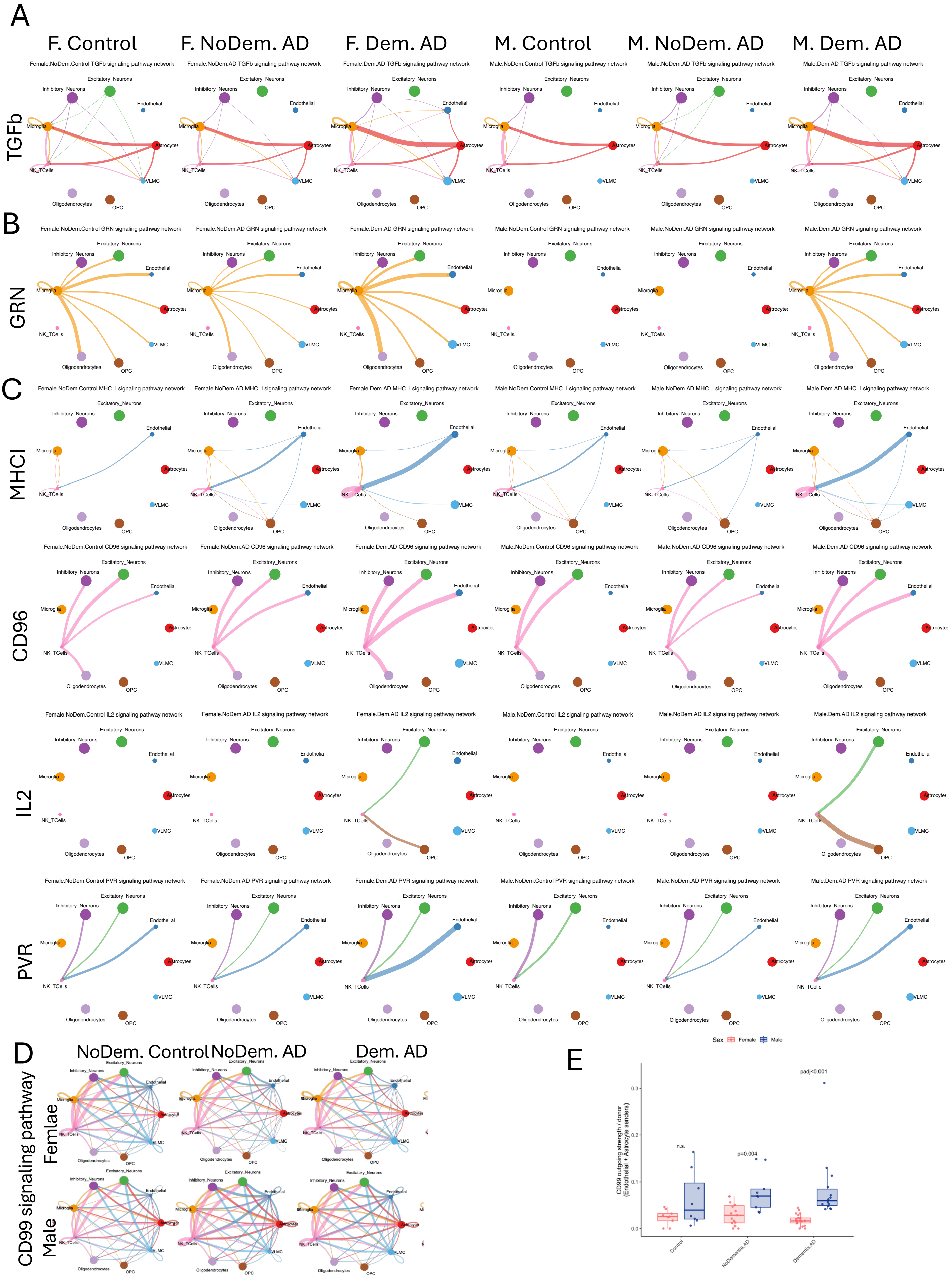

A
F. Control
F. NoDem. AD
F. Dem. AD
M. Control
M. NoDem. AD
M. Dem. AD
TGFb
Female.NoDem.Control TGFb signaling pathway network
Female.NoDem.AD TGFb signaling pathway network
Female.Dem.AD TGFb signaling pathway network
Male.NoDem.Control TGFb signaling pathway network
Male.NoDem.AD TGFb signaling pathway network
Male.Dem.AD TGFb signaling pathway network
Excitatory_Neurons
Inhibitory_Neurons
Endothelial
Microglia
Astrocytes
NK_TCells
VLMC
Oligodendrocytes
OPC
B
GRN
Female.NoDem.Control GRN signaling pathway network
Female.NoDem.AD GRN signaling pathway network
Female.Dem.AD GRN signaling pathway network
Male.NoDem.Control GRN signaling pathway network
Male.NoDem.AD GRN signaling pathway network
Male.Dem.AD GRN signaling pathway network
C
MHCI
Female.NoDem.Control MHC−I signaling pathway network
Female.NoDem.AD MHC−I signaling pathway network
Female.Dem.AD MHC−I signaling pathway network
Male.NoDem.Control MHC−I signaling pathway network
Male.NoDem.AD MHC−I signaling pathway network
Male.Dem.AD MHC−I signaling pathway network
CD96
Female.NoDem.Control CD96 signaling pathway network
Female.NoDem.AD CD96 signaling pathway network
Female.Dem.AD CD96 signaling pathway network
Male.NoDem.Control CD96 signaling pathway network
Male.NoDem.AD CD96 signaling pathway network
Male.Dem.AD CD96 signaling pathway network
IL2
Female.NoDem.Control IL2 signaling pathway network
Female.NoDem.AD IL2 signaling pathway network
Female.Dem.AD IL2 signaling pathway network
Male.NoDem.Control IL2 signaling pathway network
Male.NoDem.AD IL2 signaling pathway network
Male.Dem.AD IL2 signaling pathway network
PVR
Female.NoDem.Control PVR signaling pathway network
Female.NoDem.AD PVR signaling pathway network
Female.Dem.AD PVR signaling pathway network
Male.NoDem.Control PVR signaling pathway network
Male.NoDem.AD PVR signaling pathway network
Male.Dem.AD PVR signaling pathway network
D
NoDem. Control
NoDem. AD
Dem. AD
CD99 signaling pathway
Femlae
Male
E
Sex
Female
Male
CD99 outgoing strength / donor
(Endothelial + Astrocyte senders)
0.0
0.1
0.2
0.3
n.s.
p=0.004
padj<0.001
Control
NoDementia_AD
Dementia_AD

F

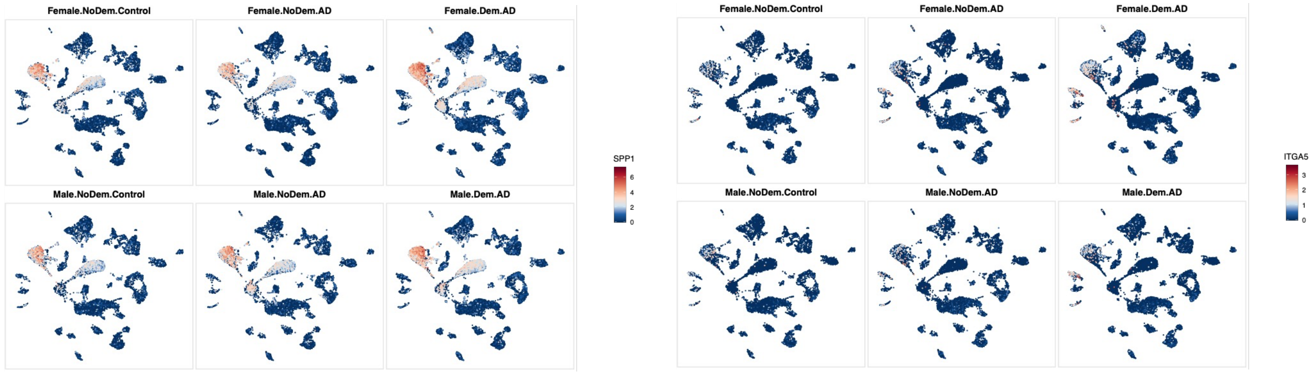


G

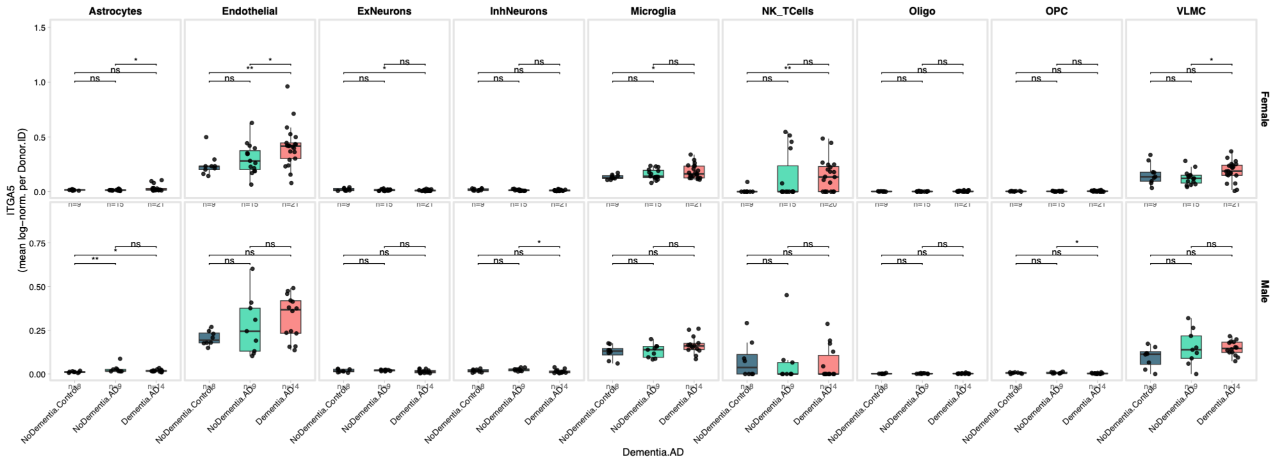


**Supplementary Figure 5. Sex-biased cell:cell signaling among microglia and other cell types in AD. A-D.** Circle plots of communication probability showing cross-condition comparison of the TGFb (A), GRN(B) and T cell incoming (C) and CD99(D) signalling pathways across 6 CellChat object(s) (p-value threshold: 0.05). **E.** Donor-level CD99 outgoing strength (Endo+Astro senders). **F.** Feature plots (UMAP) of the SPP1 ligand (left) and its integrin receptor ITGA5 (right) across the six sex-by-stage groups. SPP1 is largely restricted to microglia, whereas ITGA5 marks the endothelial and vascular compartment **G.** Pseudobulk box plot of ITGA5. Each dot represents the mean log-normalized expression for one Donor.ID (n = 76 donors), grouped by Dementia.AD, split by GlobalCellTypes, rows by Sex. Box plot elements: center line = median; box limits = 25th-75th percentile (IQR); whiskers extend to the furthest observation within 1.5x IQR from the box; outliers beyond this range are not shown. Statistical comparisons: Wilcoxon rank-sum test on donor-level means.

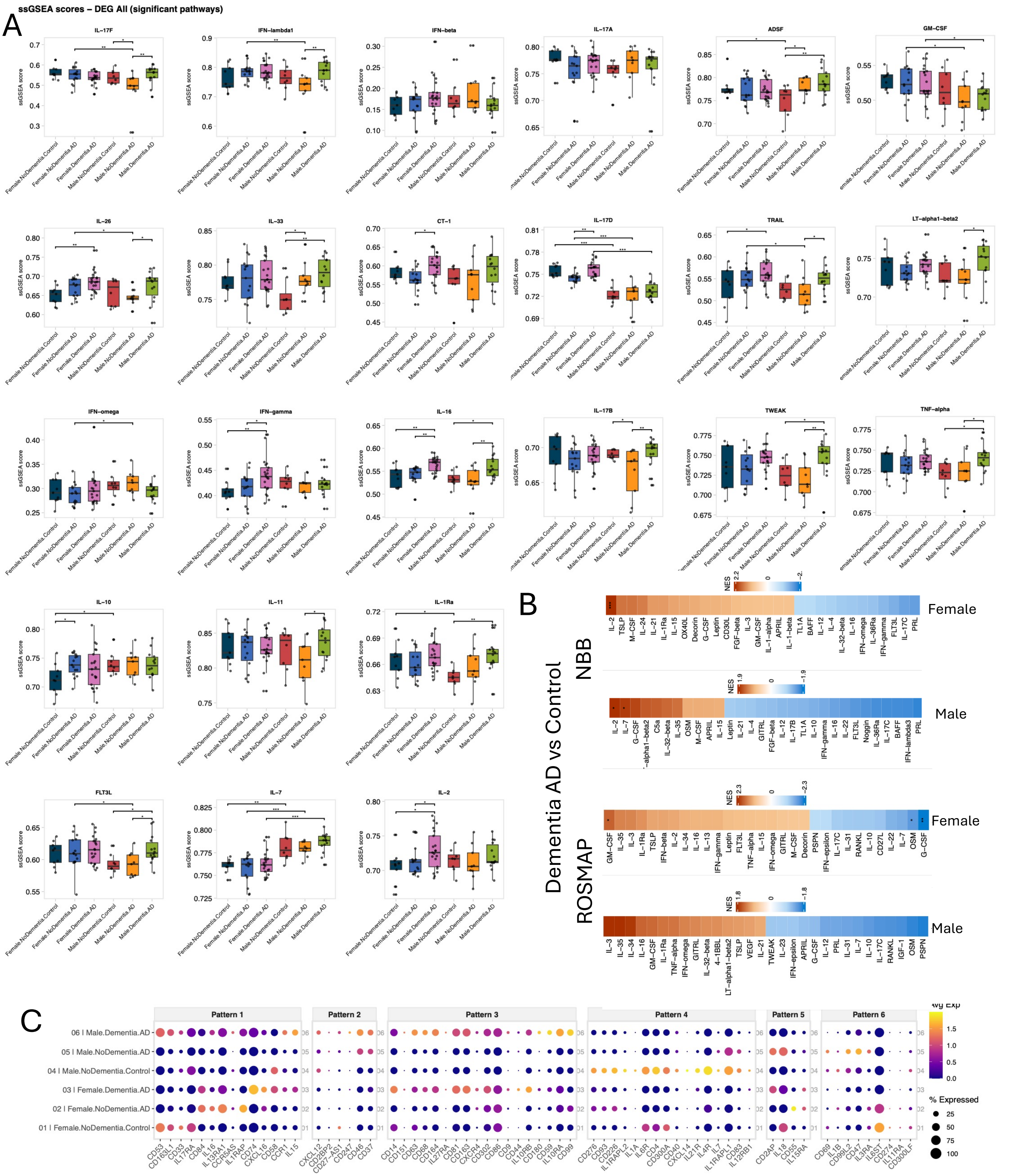


**Supplementary Figure 6. Cytokine-response signature enrichment across the six sex-by-stage groups.** A. Boxplots of single-sample GSEA (ssGSEA) enrichment scores for the cytokine-response signatures that were significantly differentially enriched across groups (Microglia; Parse cytokine gene sets). Each panel is one cytokine signature; within a panel, scores are shown per sex-by-stage group across individual donors (each point one donor), with brackets indicating significant pairwise comparisons. **B.** Cross study validation showing heatmaps of donor-level pseudobulk GSEA Normalized Enrichment Scores for the cytokine signatures, for the Dementia.AD vs NoDementia.Control, computed separately in female and male donors. Rows: top 30 signatures by max |NES|. Method: donor-level pseudobulk, edgeR TMM normalization, limma-voom moderated t-statistic pre-ranking, fgsea. NES > 0 = enriched in dementia. Stars: * BH < 0.05, ** < 0.01, *** < 0.001. **C.** Dot plot of 108 cytokine, chemokine and receptor genes (IL, CCL/CCR, CXCL/CXCR and CD gene families) across the six groups, with genes grouped into 10 clusters by hierarchical clustering. Dot size: percent of cells expressing; dot color: scaled mean expression.
